# Automated and manual segmentation of the hippocampus in human infants

**DOI:** 10.1101/2022.07.17.500316

**Authors:** J. T. Fel, C. T. Ellis, N. B. Turk-Browne

## Abstract

The hippocampus, critical for learning and memory, undergoes substantial changes early in life. Investigating the developmental trajectory of hippocampal structure and function requires an accurate method for segmenting this region from anatomical MRI scans. Although manual segmentation is regarded as the “gold standard” approach, it is laborious and subjective. This has fueled the pursuit of automated segmentation methods in adults. However, little is known about the reliability of these protocols in human infants, particularly when anatomical scan quality is low from increased head motion or shorter sequences that minimize head motion. During a task-based fMRI protocol, we collected quiet T1-weighted anatomical scans from 42 sessions with awake infants aged 4–23 months. We first had two expert tracers manually segment the hippocampus bilaterally and assess inter-rater reliability. We then attempted to predict these manual segmentations using four protocols: average adult template, average infant template, FreeSurfer software, and Automated Segmentation of Hippocampal Subfields (ASHS) software. ASHS generated the most reliable hippocampal segmentations in infants, exceeding manual inter-rater reliability of the experts. Automated methods can thus provide robust hippocampal segmentations of noisy T1-weighted infant scans, opening new possibilities for interrogating early hippocampal development.

**Highlights:**

- Inter-rater reliability of manual segmentation of infant hippocampus is moderate.
- Template-based methods and FreeSurfer provide reasonably accurate segmentations.
- ASHS produces highly accurate segmentations, exceeding manual inter-rater reliability.

## 1. Introduction

The hippocampus plays a fundamental role in learning and memory (Cohen & Eichenbaum 1993; Corkin, 2013; Nadel & Hardt, 2011; Schapiro et al., 2017; Squire, 1992; Tulving et al., 1998). This role emerges in development while the hippocampus undergoes significant changes well into adolescence (Arnold & Trojanowski, 1996; Gogtay et al., 2006; Keresztes et al., 2018; Schlichting et al., 2017; Uematsu et al., 2012). Understanding this developmental trajectory of hippocampal structure and function requires demarcating this region from surrounding regions reliably and accurately. This is perhaps most challenging in infancy when the hippocampus doubles in volume (Ellis et al., 2021a; Gómez & Edgin, 2016; Uematsu et al., 2012).

Although detailed tutorials exist to guide the segmentation of the adult hippocampus (e.g., Dalton et al., 2017), these methods are difficult to apply to the infant hippocampus. These difficulties stem from a lack of agreement about infant-appropriate anatomical landmarks that can be used to guide segmentation with magnetic resonance imaging (MRI). In adults, these landmarks were derived from histological assessments of humans (Insausti & Amaral, 2004) and non-human primates (Rakic & Nowakowski, 1981) but equivalent landmarks have generally not been established in infancy (cf. Insausti et al., 2010). Additionally, infants tend to move their heads more during MRI, even while asleep (Denisova, 2019). This movement can blur boundaries between white and gray matter that define the hippocampus. Furthermore, sound levels limit the use of certain sequences in minimal risk research. For example, T2-weighted scans — typically acquired for adult hippocampal segmentation — can wake or startle sleeping infants. Conversely, T1-weighted scans can be quieter, but typically have a voxel resolution around 1 mm (compared to 0.4−0.5 mm from a T2-weighted scan), obscuring landmarks needed to identify the hippocampus. Finally, the hydration properties of myelin change rapidly over the first year of life, altering the contrast of white matter in T1-weighted images (Xue et al., 2007) and lowering tissue contrast (Shi et al., 2011). For these reasons, segmentation of the infant hippocampus faces several unique challenges not shared with hippocampal segmentation in adults or even older children. These limitations have led to a focus on gross segmentation of the whole infant hippocampus (Gousias et al., 2012; Guo et al., 2014; Guo et al., 2015; Ellis et al., 2021a), although there are recent attempts to segment subfields (Li et al., 2019; Zhu et al., 2019).

Manual segmentation is often considered the “gold standard” for accurately defining regions of interest (ROI) like the hippocampus (Boccardi et al., 2015; Frisoni et al., 2015; Rodionov et al., 2009). Protocols for manually segmenting the whole infant hippocampus have been developed (Gousias et al., 2012). However, manual segmentation requires considerable training and is time-consuming even for experts (Morey et al., 2009), and the results can suffer from subjective biases (Colon-Perez et al., 2016). Automated methods can be more efficient (Morey et al., 2009) and reduce bias (Khlif et al., 2019). Given a sample of adult T2-weighted and T1-weighted MRI scans, multi-atlas automated algorithms such as Automated Segmentation of Hippocampal Subfields (ASHS; Xie et al., 2019; Yushkevich et al., 2015) generate accurate and reliable hippocampal subfield segmentations from a library of manually labeled MRI scans. FreeSurfer (Brown et al., 2020; Iglesias et al., 2015) employs a statistical atlas from high resolution (~0.1 mm) *ex vivo* MRI scans for adult hippocampal segmentation. Tested on child MRI data (> 6 years), ASHS produces consistently accurate segmentations (Bender et al., 2018; Schlichting et al., 2019), whereas FreeSurfer can fail to approximate manual segmentations (Schoemaker et al., 2016). Although the efficacy of these algorithms for adult and child data is well known, their accuracy and reliability in infants is still emerging.

Automated approaches such as a dilated dense network embedded within a U-net (Zhu et al., 2019) and a multi-atlas-based protocol (Guo et al., 2015) can generate reliable segmentations of the infant hippocampus using T1-weighted scans. The metric for evaluating reliability in these previous studies was *intra-rater* reliability — how accurately the model predicts a held-out manual segmentation from the same expert tracer who created the other segmentations on which the model was trained (i.e., training on expert A, testing on expert A). However, this metric cannot distinguish whether the algorithm has learned the idiosyncrasies of this tracer or more generalized segmentation principles, a version of the overfitting problem in machine learning (Carmo et al., 2019). To determine whether automated methods create segmentations that generalize, it is necessary to employ *inter*-rater reliability (IRR) as the performance metric — how accurately the model predicts a manual segmentation from an expert tracer whose segmentations were held out of model training entirely (i.e., training on expert A, testing on expert B). This metric is particularly important in cases where subjectivity has a strong influence, such as when the scans used for segmentation have high noise.

In the present study, we used this more stringent across-expert IRR test of generalizability to investigate four protocols for infant hippocampal segmentation. As tested in older children (Schlichting et al., 2019), we evaluated how well the manual segmentation of an infant’s hippocampus could be predicted from: (1) manual segmentations of adults (average adult template), (2) manual segmentations of other infants (average infant template), (3) automated FreeSurfer segmentation of the same infant, and (4) automated ASHS segmentation of the same infant. For each infant, two independent expert tracers manually segmented the whole hippocampus and medial temporal lobe (MTL) cortex, enabling a fully-manual benchmark measurement of IRR against which to evaluate the protocols (Bender et al., 2018). As a further test of robustness, these protocols were applied to relatively low-quality data: short T1-weighted scans collected from 42 sessions with infants aged 4–23 months who were awake in the scanner. This provides a lower bound on data quality that might be expected from infant data collection during sleep or sedation, such that success may bode well for broader application, including to lower-quality data from portable scanners (Deoni et al., 2021). Success with awake scans is also important because infant fMRI is becoming more feasible, which could enable task-based studies of hippocampal function during online learning and memory (Ellis et al., 2021a).

## 2. Methods

### 2.1. Participants

Data from 42 sessions with 22 distinct healthy infants (13 female) were collected for this study. At the time of each session, the infants ranged in age from 3.6 to 23.1 months (M=10.6, SD=5.1). These sessions were held at three sites: 7 at the Scully Center for the Neuroscience of Mind and Behavior at Princeton University, 10 at the Magnetic Resonance Research Center (MRRC) at Yale University, and 25 at the Brain Imaging Center (BIC) at Yale University. The data were collected as part of ongoing task-based fMRI studies in awake infants. These sessions were chosen for segmentation because these participants completed a study that examined hippocampal contributions to infant learning (e.g., Ellis et al., 2021a). Of the 42 anatomical scans from these sessions, 10 were from “non-repeat” infants who contributed one session; “repeat” infants contributed more than one session: 6 had two sessions (12 total), 5 had three sessions (15 total), and 1 had five sessions. At least one month passed between repeat scans (range: 1.1–6.4). Refer to Table S2 for information on each participant. Parents gave informed consent on behalf of their child to a protocol approved by the Institutional Review Board at each site.

### 2.2. Data Acquisition

MRI images were acquired with a Siemens Skyra (3T) MRI scanner at Princeton and a Siemens 3T Prisma MRI scanner at both the MRRC and BIC sites at Yale. The bottom half of the 20-channel Siemens head coil was used at all sites. The top half of the head coil was removed during these studies to permit visual stimulation and eye-tracking, and to monitor infant comfort and task compliance. We previously validated this coil configuration, including showing similar signal from the bottom half alone in infants as the top and bottom in adults, given the smaller head size of infants (Ellis et al., 2020). Regardless, these scans provide a conservative test of the segmentation algorithms, and they are representative of our approach to awake, task-based infant fMRI (Ellis et al., 2021a, 2021b, 2021c). Infant anatomical images were acquired with a T1-weighted PETRA quiet sequence (TR1=3.32ms, TR2=2250ms, TE=0.07ms, flip angle=6°, matrix=320×320, slices=320, resolution=0.94mm iso, radial slices=30000, duration= 3mins, 8s).

### 2.3. Procedure

Our protocol for collecting infant MRI data has been described in detail in a previous methods paper (Ellis et al., 2020). Briefly, parents and infants were acclimated to the scanning environment during an orientation visit prior to their first scanning session. Parents scheduled sessions for the time of day they felt the infant would be calmest and happiest. Both the parent and infant were rigorously screened for metal. Hearing protection was applied to the infant in three layers: silicon inner ear putty, over-ear adhesive covers, and earmuffs. The infant was carefully placed on a vacuum pillow on the scanner bed, comfortably reducing movement. During the anatomical scans, infants were shown a movie to keep them engaged or they completed an eye-tracking calibration. If the infant became fussy, they were shown something else until they calmed down, or we ended the scan for a break before re-trying. In some cases (N=8), we initiated an anatomical scan because the infant fell asleep after being awake for other parts of the study. Infants were monitored during data collection with an in-bore MRC video-camera. If the infant moved excessively, the anatomical scan was stopped to minimize wasted time in producing a blurry image. In some sessions, multiple anatomical scans were collected. The highest quality scan was chosen for segmentation. When two high-quality scans were acquired, they were aligned and averaged (N=10).

### 2.4. Transformation to Standard Space

For the comparison of scans across sessions and for the construction of an average infant template, infant data were aligned into standard space using nonlinear methods (see Figure S1 for linear methods). Past research has found that nonlinear methods result in more accurate alignment of an individual infant’s anatomical scan to an infant template (Pineda et al., 2014). The nonlinear method we used was ANTs, based on diffeomorphic symmetric normalization (Avants et al., 2011). After aligning each scan to an age-specific MNI infant template (Fonov et al., 2011) with ANTs, we further aligned the data from this infant template space to an adult MNI template (MNI152; Mazziotta et al., 2001) using a precomputed linear mapping.

### 2.5. Manual Segmentation

Two tracers, CE and JF (authors), manually segmented all 42 anatomical scans in accordance with an infant hippocampal segmentation protocol (Gousias et al., 2012) supplemented by guidance from an adult MTL cortex demarcation protocol (Aly & Turk-Browne, 2016). The tracers were blind to the age and sex of the participants, as well as to the corresponding manual segmentations of the other tracer.

The infant hippocampus is more easily visible using sagittal slices than coronal slices (Gousias et al., 2012). The lateral edge of the hippocampus was defined on the sagittal plane by the white matter of the temporal lobe and the grey matter of the hippocampus, although on some occasions cerebrospinal fluid (CSF) was used to define this edge and not white matter. The medial edge was defined using the lateral ventricle. The amygdala helped discriminate the anterior boundary for the head of the hippocampus, especially in more lateral sagittal slices. However, traversing medially, it was common for this anterior edge distinction to become less clear. In these instances, the hippocampal shape from adjacent slices guided the estimate of the anterior edge. The posterior edge at the tail of the hippocampus was generally circumscribed by white matter or CSF. On more medial slices, it was common for the body to disappear, with only the posterior-tail and anterior-head visible. After the hippocampus was fully traced in the sagittal plane, both tracers refined the outlines using the coronal plane, ensuring that the subiculum extended medially in this coronal view and did so consistently across sagittal slices.

MTL cortex, encompassing entorhinal, perirhinal and parahippocampal cortices, was segmented on the coronal plane. As with adult MTL cortex (Aly & Turk-Browne, 2016), the anterior extent of infant MTL cortex was determined by the collateral sulcus (CS). The CS was followed from anterior to posterior coronal slices and MTL cortex was drawn as connecting to the hippocampus. In some scans, the CS was inconsistently visible across slices. When the CS was not visible, tracers interpolated between landmarks and drew MTL cortex despite uncertainty. The posterior extent of MTL cortex was determined by an imaginary line perpendicular to the hippocampal body.

### 2.6 Dice Evaluation

The Dice metric (Dice, 1945) was used throughout to make comparisons between segmentations of regions. Dice measures the spatial overlap between two volumes, calculated as:

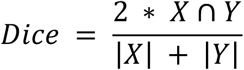

where *X* and *Y* denote the set of labeled voxels to be compared. The value of a Dice metric ranges from 0, which indicates no spatial overlap between the two sets of manual segmentations, to 1, indicating perfect overlap. Values below 0.5 were considered weak, values between 0.5–0.6 were considered moderate, values between 0.6–0.7 were considered satisfactory, and values above 0.7 were considered robust (Hashempour et al., 2019). To compute IRR, the Dice between CE and JF’s nonlinearly aligned segmentations of a given scan was calculated separately for the hippocampus and MTL cortex and this was repeated for all 42 scans. These calculations were performed in standard space in order to make the comparisons consistent across use cases; however, IRR values were highly similar in native anatomical space. Dice was used for other comparisons as well, as described below.

### 2.7. Repeat Analyses

Repeat scans from the same infant across sessions afforded the opportunity to investigate whether the location of the hippocampus in the developmental future or past could be predicted from a segmentation at a younger or older age, respectively. As a control, each pair of repeat sessions from the same infant was matched in age to two sessions from different infants as closely as possible (error in age matching: M=0.31 months, SD=0.38, range=0.00–1.40). The Dice of repeat sessions within-participant was compared with the Dice of one of those repeat sessions with the control session age-matched to the other repeat session in the pair (and then reversed to control for the other repeat session). For example, consider the comparison between the first session of participant s7017_1_1 (5.2 months old) and their fourth session s7017_1_4 (8.3 months old). An age-match control for the first session was found with participant s0687_1_1 (5.3 months old), with an age match error of 0.1 months. An age-match control for the latter session was s8607_1_1 (8.5 months old), with an age match error of 0.2 months. We thus compared s7017_1_1/s7017_1_4 (within participant) vs. s7017_1_1/s8607_1_1 and s0687_1_1/s7017_1_4 (between participants). The whole procedure was then run for all pairings of repeat sessions.

### 2.8. Average Adult Template

An average adult template was constructed using the Harvard-Oxford subcortical probabilistic atlas for the hippocampus (Makris et al., 2006). This probabilistic atlas was originally created by aggregating and averaging the binarized hippocampal segmentations from 37 distinct healthy adults (ages 18-50). Hence, each voxel value reported the proportion of participants for whom that voxel was labeled as hippocampus. In the present study, this atlas was thresholded at a probability of 50% and binarized to create our average adult template. This was only done for the hippocampus since no MTL cortical regions were segmented in this atlas. Dice was then used to compare this template with the nonlinearly aligned infant manual segmentations from CE and JF.

### 2.9. Average Infant Template

To produce an average infant template, a leave-one-scan-out approach was employed. On each iteration, all segmentations from the two manual tracers were aggregated except those of the held-out scan. The aggregated segmentations were averaged and thresholded, requiring voxels to be labeled in at least 50% of sessions to be included. Dice was then used to compare this template to each tracer’s segmentation of the held-out scan.

### 2.10. FreeSurfer

FreeSurfer served as an automated algorithm. Segmented infant hippocampal data (no MTL cortex available) were extracted using an adult FreeSurfer reconstruction pipeline (Iglesias et al., 2015). In some sessions, FreeSurfer failed to identify the hippocampus. In such cases, FreeSurfer was applied to other versions of the same scan (e.g., with skull stripping). With this procedure, FreeSurfer succeeded on 40 of 42 scans. The two scans that FreeSurfer was unable to segment were excluded from the calculation of the FreeSurfer IRR metric. This metric was calculated by comparing the FreeSurfer segmentation of an infant to each tracer’s corresponding manual segmentation.

### 2.11. ASHS

Multiple ASHS models were evaluated in this study. The “Adult-Pretrained-ASHS” model was previously trained by the Penn Image Computing and Science Laboratory (PICSL) to automatically segment the adult hippocampus and MTL (Xie et al., 2016). Importantly, PICSL used only T1-weighted scans from adults aged 55 and older for model training. Given that we segmented the whole hippocampus in the present study, demarcations of subregions produced by this model (e.g., anterior and posterior hippocampus) were pooled.

The other ASHS models were trained on our manual infant segmentations aligned into standard space. We initially attempted to supply ASHS with the segmentations in each infant’s native anatomical space; however, the ASHS pipeline failed to align these data appropriately when forming the template. Supplying the segmentations in standard space, based on brains aligned with an optimized nonlinear approach, reduced the burden on the alignment step within ASHS. We made additional modifications to the ASHS configuration file to increase the search iterations (refer to the data release for the updated parameters).

A cross-validation approach was adopted for these training sessions, in which an ASHS model was trained on all but one scan. The model was then used to predict the nonlinearly aligned segmentation of the held-out scan. The “CE-ASHS” and “JF-ASHS” models were trained using the manual infant segmentations from CE and JF manual tracers, respectively. The “Infant-Trained-ASHS” model was trained over *both* CE and JF’s manual infant segmentations. On each leave-one-scan-out training iteration for this latter model, the order of CE and JF’s segmentations was shuffled randomly to prevent order effects (Mange, 2019).

In addition to this leave-one-scan-out approach, we performed an even more stringent test of generalization by omitting all scans from one participant from the training (some participants had more than one scan). These leave-one-participant-out (LOPO) versions of the models are “LOPO-CE-ASHS”, “LOPO-JF-ASHS”, and “LOPO-Infant-Trained-ASHS”, respectively (see Table S1; Figure S5). The automated segmentation of the left-out-participant scan generated from each ASHS model was evaluated against both tracers’ manual segmentations of that same left-out-participant scan, resulting in two Dice measurements.

### 2.12. Volume Bias

To further evaluate FreeSurfer and ASHS, we examined the bias present in the set of automated segmentations from each algorithm (Schlichting et al., 2019). Bland-Altman plots (Bland & Altman, 2007) were used to quantify the extent to which the volume of the hippocampus was over- or under-estimated by these two algorithms. The volumes of the manually segmented hippocampus from the two tracers were averaged and the segmentations from each automated method were used for comparison. For each participant, total hippocampal volume was calculated by summing the manual volume (averaged across tracers) with the corresponding automated volume:

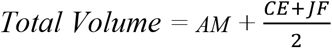

Where *AM, CE,* and *JF* denote the number of labeled hippocampal voxels from an automated method and tracers CE and JF, respectively. This was then plotted against the difference in hippocampal volume between the automated method and the manual average:

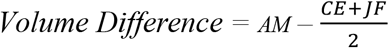

If an automated method has bias, then this would be reflected in volume difference values above or below zero (i.e., over- or under-estimation). Bias may vary with total volume, which would manifest as a statistically significant relationship between volume and error (i.e., the model shows different bias across a range of hippocampal volumes).

### 2.13. Statistics

A non-parametric bootstrap resampling approach was used to determine confidence intervals and statistical significance (Efron & Tibshirani, 1986). For a given measure of interest, the values across participants were first averaged. A sampling distribution of this test statistic was estimated by sampling the same number of values with replacement 10,000 times and averaging on each iteration. Confidence intervals were determined based on the percentiles of this sampling distribution. Statistical significance was determined by the proportion of iterations whose mean had the opposite sign from the true effect, doubled to make the p-value two-tailed. A similar bootstrap resampling procedure was used to evaluate correlations, sampling bivariate data with replacement 10,000 times and calculating the Pearson correlation on each iteration. The p-value was calculated as the proportion of samples with a correlation coefficient of the opposite sign from the true correlation, doubled to make the test two-tailed.

### 2.14. Data and Code Availability

Data will be made available upon publication of this manuscript. The code for performing the reported analyses can be found here: https://github.com/ntblab/infant_neuropipe/tree/MTL_Segmentation/scripts/MTL_Segmentation.

## 3. Results

### 3.1. Inter-Rater Reliability

The average Dice between tracers across the 42 sessions was 0.57 (SD=0.07, range=0.43–0.70). This moderate IRR reflects the challenging nature of segmenting infant MRI data. Figure 1 depicts an above-average-quality and below-average-quality infant T1-weighted MRI scan in anatomical space. Examining the above-average-quality scan, the tracers show high agreement in both lateral slices (Figure 1A) and medial slices (Figure 1B). For the below-average-quality scan, while the tracers show high agreement on some slices (Figure 1C), they do not show the same level of agreement on others (Figure 1D). The disagreement about placement of the hippocampal head arises from low contrast tissue in this scan and the ambiguity of the hippocampus/amygdala boundary in the presence of MRI noise. Nevertheless, by traditional methods that treat manual segmentation as the “gold standard”, IRR represents a rigorous benchmark against which the four protocols below can be evaluated. Specifically, the IRR Dice reflects the extent to which a given participant’s manual segmentation from one expert can be predicted from another expert, which can be fairly compared against the prediction made by each of the protocols.

**Figure 1:**
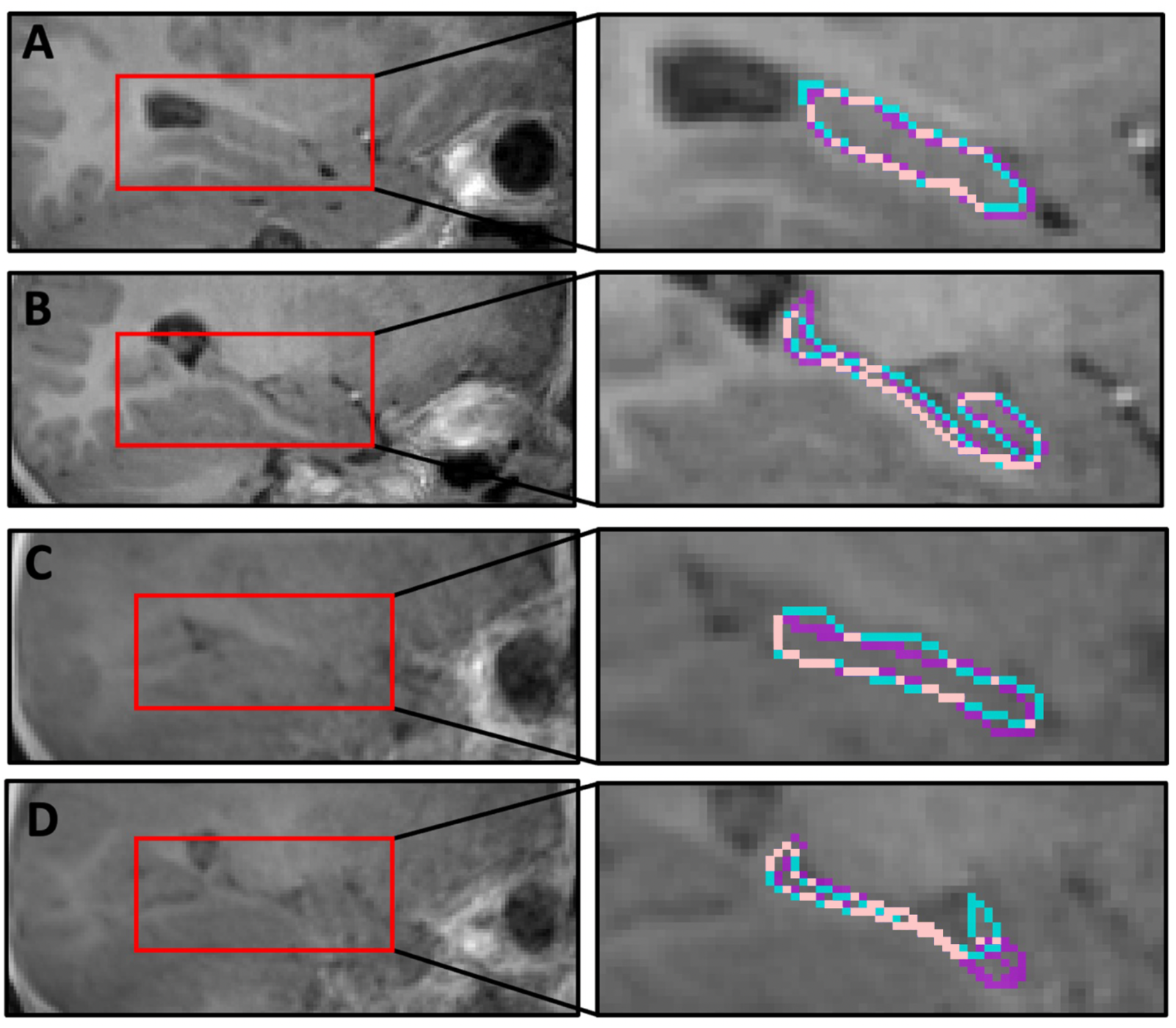
Tracing an above-average-quality and below-average-quality T1-weighted scan. The traced outline of the hippocampus on all four slices is shown in color (CE=blue, JF=purple, overlap=pink). The top two slices (above-average-quality) come from participant s2307_1_1 (19.9 months) and had a Dice of 0.63. There is agreement between CE and JF in (A) a lateral sagittal slice and (B) a middle sagittal slice. The bottom two slices (below-average-quality) come from participant s6057_1_2 (7.6 months) and had a Dice of 0.51. There is again agreement between CE and JF in (C) a lateral sagittal slice, two slices after the hippocampus first emerges in this sagittal view. However, there is disagreement about the placement of the hippocampal head in (D) a middle sagittal slice.

### 3.2. Repeat Participants

Twelve participants were scanned in more than one session, allowing Dice comparisons from the same participant within tracer (Figure S2A). Despite developmental changes in the intervening time (DeMaster et al., 2014), this repeat analysis can serve as one proxy for test-retest reliability. That is, it quantifies the degree to which the tracer reproduces their traces when segmenting similar anatomy. The average repeat Dice after nonlinear alignment to standard space was 0.60 for CE (SD=0.08, range=0.45–0.76) and 0.60 for JF (SD=0.07, range=0.43–0.72). As a control, we calculated Dice between these segmentations and segmentations from other participants matched in age. The resulting average Dice was lower for both CE (Dice=0.51, SD=0.07, range=0.28–0.63) and JF (Dice=0.47, SD=0.07, range=0.28–0.60).

### 3.3. Average Adult Template Analyses

An average adult template, constructed using the Harvard-Oxford atlas, was evaluated in terms of how well it predicted the nonlinearly aligned manual segmentations (Figure 2). The average Dice between this template and the infant hippocampal data of both tracers (CE: Dice=0.42, SD=0.09, range=0.26–0.58; JF: Dice=0.47, SD=0.07, range=0.30–0.60) was significantly lower than the manual IRR of 0.57 (CE: M=−0.15, CI=[−0.179, −0.115],*p*<0.001; JF: M=−0.10, CI=[−0.130, −0.074], *p*<0.001). There were no significant correlations between age and the success of the average adult template in predicting segmentations from either tracer (Figure S2B).

**Figure 2:**
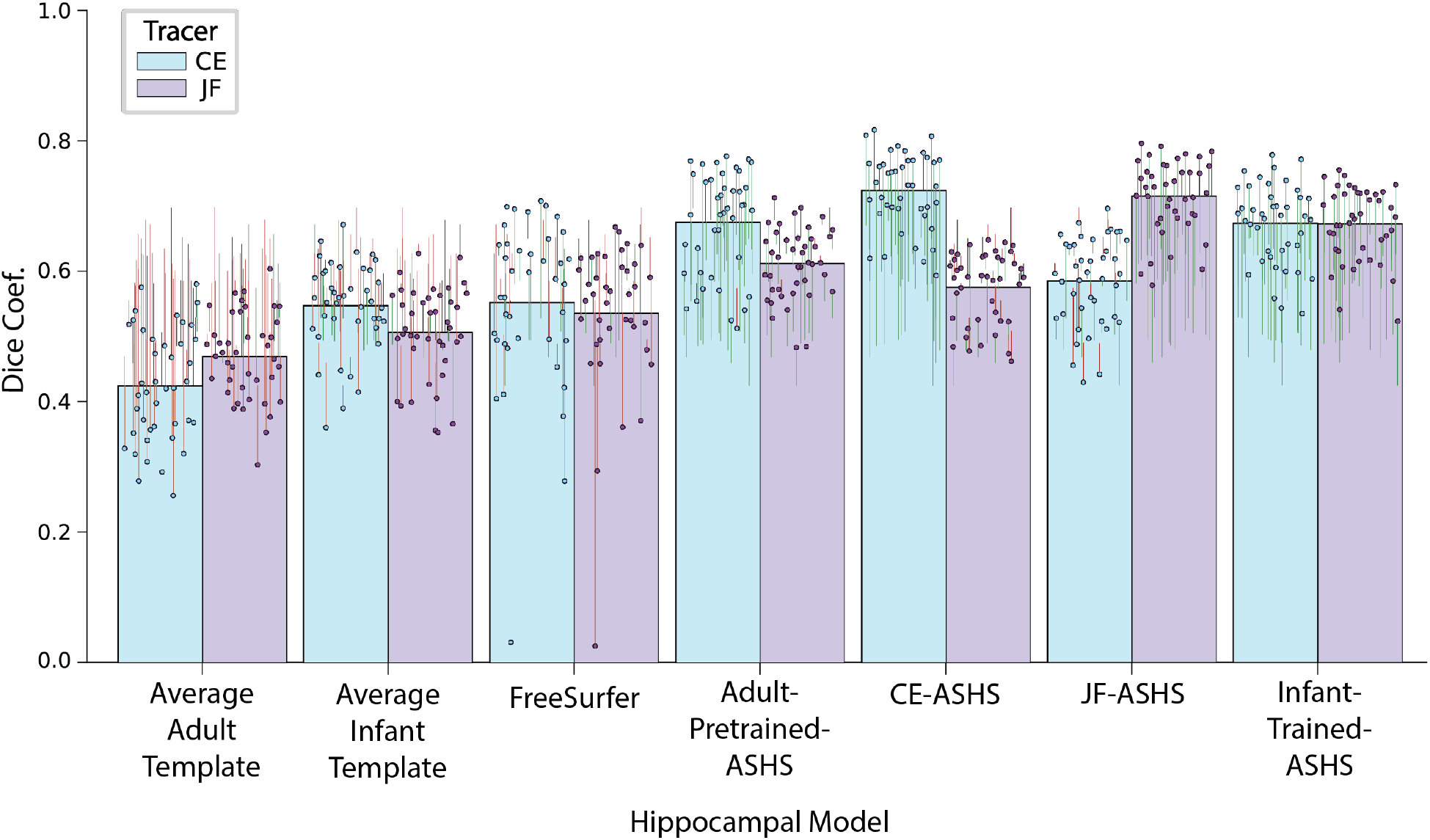
Prediction of manual segmentations of infant hippocampus by average templates, FreeSurfer, and ASHS models. The manual infant segmentations from CE and JF were nonlinearly aligned into standard space and predicted from the average adult segmentation (Average Adult Template) and the average of segmentations from other infants (Average Infant Template). The manual infant segmentations from CE and JF in subject space predicted by an automated FreeSurfer segmentation (FreeSurfer). The manual infant segmentations from CE and JF in standard space predicted by ASHS models trained on an adult library (Adult-Pretrained-ASHS), CE’s infant segmentations (CE-ASHS), JF’s infant segmentations (JF-ASHS) and both CE and JF’s infant segmentations (Infant-Trained-ASHS). The height of the colored bar indicates the average Dice for the corresponding protocol, predicting the segmentation by CE (blue) and JF (purple). Each colored dot is the Dice under that protocol for one infant session. The line extending from the dot indicates how much the protocol improves (green) or worsens (red) performance relative to the between-tracer IRR for that session.

### 3.4 Average Infant Template Analyses

We assessed an average infant template, constructed by aggregating infant segmentations, in terms of how well it predicted the manual hippocampal segmentations of a left-out session (Figure 2). This average infant template was able to adequately approximate the left-out infant data from both tracers (CE: Dice=0.55, SD=0.07, range=0.36–0.67; JF: Dice=0.51, SD=0.07, range=0.35–0.63). The metric for CE was only marginally lower than manual IRR (M=−0.02, CI=[−0.048, 0.001], p=0.064), though was significantly lower for JF (M=−0.06, CI=[−0.087, −0.043], p<0.001). Age was positively correlated with the similarity to these templates for one tracer but not the other (Figure S2C). These results suggest that nonlinear alignment methods allow for the construction of an average infant template that comes close to approximating IRR.

### 3.5. FreeSurfer Analyses

The first automated method we considered was FreeSurfer. Although not designed to work with low grey/white matter contrast, we included FreeSurfer because it is a popular method in adults (Figure 2). Infant segmentations from FreeSurfer did a reasonable job at predicting the manual infant segmentations from both tracers (CE: Dice=0.55, SD=0.13, range=0.03–0.71; JF: Dice=0.54, SD=0.12, range=0.03–0.67). The metric from CE was similar to manual IRR (M=−0.02, CI=[−0.060, 0.013], *p* = 0.283), though was significantly lower for JF (M=−0.04, CI=[−0.075, −0.007], *p*=0.013). The accuracy of FreeSurfer prediction of manual segmentations from both tracers correlated positively with participant age (Figure S3A). Note that FreeSurfer completely misplaced the hippocampus in one participant, leading to a Dice of 0.03 for both tracers (Table S3). Despite this outlier, FreeSurfer provided accurate infant hippocampal segmentations that predicted manual segmentations.

### 3.6. ASHS Analyses

The second automated method we considered was ASHS. The Adult-Pretrained-ASHS was used as a baseline for ASHS performance because it was not trained on any infant data. Notably, the average Dice between the infant segmentations from this model and the manual segmentations were robust (CE: Dice=0.68, SD=0.08, range=0.51–0.78; JF: Dice=0.61, SD=0.05, range=0.48– 0.71). These metrics significantly surpassed IRR for both tracers (CE: M=0.10, CI=[0.086, 0.123], *p*<0.001; JF: M=0.04, CI=[0.030, 0.052], *p*<0.001). The accuracy of Adult-Pretrained-ASHS prediction of manual segmentations from both tracers correlated positively with participant age (Figure S3B).

We then trained several ASHS models on infant rather than adult data, with the goal of further improving prediction by tailoring the model training to the infant brain. In previous studies (Bender et al., 2018, Schlichting et al., 2019; Yushkevich et al., 2015), ASHS could predict out-of-sample child or adult segmentations from the tracer on which it was trained. This may work in infants too if the scans contain consistent features across participants. We thus first trained separate ASHS models for each of the two expert tracers (CE-ASHS and JF-ASHS). Both models produced automated hippocampal segmentations that predicted a held-out manual segmentation from the tracer on which they were trained extremely well (Table S4; CE-ASHS predicting CE: Dice=0.73, SD=0.06, range=0.59–0.82; JF-ASHS predicting JF: Dice=0.72, SD=0.06, range=0.58–0.80). These metrics significantly exceeded IRR, an effect found in every participant (CE-ASHS predicting CE: M=0.15, CI=[0.137, 0.172], *p*<0.001; JF-ASHS predicting JF: M=0.14, CI=[0.129, 0.159], *p*<0.001). Segmentation accuracy for both models correlated positively with participant age (Figure S3C, D).

The ability of the models above to predict segmentations from the same tracer may reflect idiosyncrasies of that tracer’s approach, a form of overfitting. We thus applied these same models to segmentations from the other tracer (to which the models were blind during training). This provides as a more conservative test of how well the models generalize. Both models were able to predict the other tracer’s manual segmentations (Table S4; JF-ASHS predicting CE: Dice=0.59, SD=0.07, range=0.43–0.70; CE-ASHS predicting JF: Dice=0.58, SD=0.05, range=0.46–0.65). These metrics were similar to manual IRR (JF-ASHS predicting CE: M=0.01, CI=[−0.002, 0.030], *p*=0.088; CE-ASHS predicting JF: M=0.00, CI=[−0.008, 0.019], *p*=0.512). In addition, CE-ASHS and JF-ASHS automated prediction of manual JF and CE hippocampal segmentations, respectively, correlated positively with participant age (Figure S3C, D). Altogether, ASHS models trained on just one tracer can generate segmentations that capture what is shared between tracers.

We further trained a combined infant ASHS model jointly on segmentations from both tracers and tested it on held-out manual segmentations from each tracer. This automated Infant-Trained-ASHS model predicted manual infant segmentations quite well (Table S4; predicting CE: Dice=0.67, SD=0.06, range=0.54–0.78; predicting JF: Dice=0.67; SD=0.06, range=0.52–0.76) and better than IRR (CE: M=0.10, CI=[0.086, 0.119], *p*<0.001; JF: M=0.10, CI=[0.090, 0.113], *p*<0.001). Additionally, the accuracy of this model correlated positively with participant age for both tracers (Figure S3E). Of note, the ASHS model trained with both tracers was less reliable than the single-tracer models predicting the manual segmentations of the same tracer (Infant-Trained-ASHS vs. CE-ASHS predicting CE: M=−0.05, CI=[−0.062, −0.041], *p*<0.001; JF-ASHS predicting JF: M=−0.04, CI=[−0.052, −0.034], *p*<0.001). However, the combined model significantly outperformed single-tracer model predictions of the manual segmentations from the other tracer (Infant-Trained-ASHS vs. JF-ASHS predicting CE: M=0.09, CI=[0.080, 0.097], *p*<0.001; CE-ASHS predicting JF: M=0.10, CI=[0.087, 0.107], *p*<0.001).

We sought to understand why the ASHS models trained on one tracer could accurately predict segmentations from the other tracer, as well as why the combined Infant-Trained-ASHS model better captured what was shared between tracers. We examined the degree to which the segmentations generated from these three models matched an optimal representation of what was shared across tracers — an intersection of the manual segmentations from CE and JF. This was considered optimal because every voxel in the intersection is, by definition, guaranteed to match between tracers, ensuring a maximally high IRR while still maintaining a high within-tracer reliability. After calculating the average Dice between these intersections and each tracer’s set of manual segmentations (CE: Dice=0.74, SD=0.06, range=0.59–0.84; JF: Dice=0.72, SD=0.07, range=0.57–0.84), an “optimal Dice” was created by averaging across tracers (Dice=0.73, SD=0.06, range=0.60−0.82). ASHS models trained with one tracer’s data achieved 82% of the optimal Dice (CE-ASHS: Dice=0.60, SD=0.06, range=0.48–0.70; JF-ASHS: Dice=0.60, SD=0.09; range=0.43–0.74). The ASHS model trained on both tracers achieved 88% of the optimal Dice (Dice=0.64, SD=0.09, range=0.43–0.74).

### 3.7. MTL Analyses

We investigated whether the pattern of results observed with the hippocampus would generalize to segmentations of the MTL cortex. The average IRR across participants for MTL cortex, aligned to standard space, was 0.52 (SD=0.09, range=0.32–0.70). A nonlinearly aligned infant template was able to approximate IRR for both tracers (Figure S4), similar to the hippocampus. Given that no MTL cortex mask was available for use in the Harvard-Oxford subcortical template (Makris et al., 2006), no MTL cortex analyses were able to be run with the average adult template. The CE-ASHS and JF-ASHS models produced MTL segmentations that predicted manual segmentations within and between tracers more reliably than IRR (Figure S4), with the Infant-Trained-ASHS model exceeding IRR for both tracers as well (Figure S4). These data validate ASHS as a general tool for segmenting different regions throughout the infant brain. The FreeSurfer reconstruction pipeline employed in this study (Iglesias et al., 2015) had no MTL cortex segmentation available.

### 3.8. Volume Bias

To further evaluate the efficacy of the automated methods, we measured bias in hippocampal volume using Bland-Altman plots (Figure 3; Bland & Altman, 2007). These scatterplots display the total volume of a region against the difference in size of the regions between manual and automatic segmentations. Bias can manifest in different ways: the model may only be accurate for participants with a small hippocampus and either under-estimate or over-estimate larger hippocampal volumes, resulting in a negative or positive correlation, respectively. Or the model may only be accurate for participants with a large hippocampus and either under-estimate or over-estimate smaller hippocampal volumes, resulting in a positive or negative correlation, respectively. If the automated model consistently over or under-estimates a region size, the predictions will all be above or below zero, respectively.

**Figure 3:**
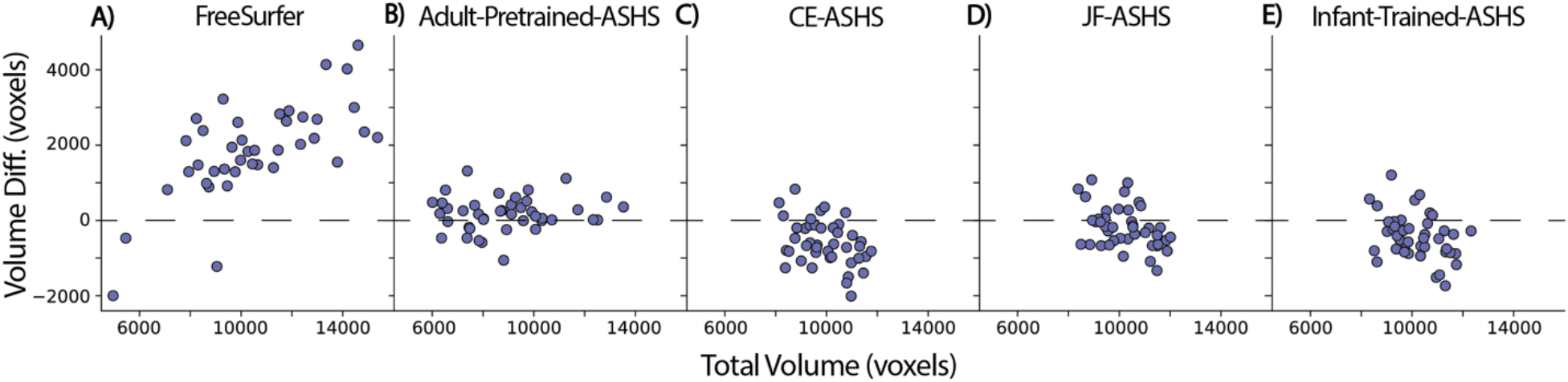
Calculating bias for each automated algorithm. Bland-Altman plots of volume difference estimation between manual segmentation and (A) FreeSurfer, (B) Adult-Pretrained-ASHS, (C) CE-ASHS, (D) JF-ASHS, or (E) Infant-Trained-ASHS. For each bias plot, the x-axis represents the average volumes of the manually segmented hippocampus of the two tracers summed with the automated volumes; the y-axis represents the difference between the automated and average manual volumes; the dashed axis lines represent a volume difference of zero.

FreeSurfer shows clear evidence of bias (Figure 3A). The mean difference between FreeSurfer segmentations and the manual segmentations of CE and JF (M=1881 voxels, CI=[1484, 2262], *p*<0.001) revealed that FreeSurfer considerably over-estimated hippocampal volume, as has been shown previously (Schmidt et al., 2018; Schoemaker et al., 2016). FreeSurfer also shows a greater bias for larger hippocampal volumes (*r*=0.68, *p*<0.001). The ASHS segmentations also differed in overall volume from the manual segmentations of CE and JF, with over-estimation by Adult-Pretrained-ASHS (Figure 3B; M=176 voxels, CI=[41, 310], *p*==0.010), and underestimation by CE-ASHS (Figure 3C; M=−544 voxels, CI=[−728, −364], *p*<0.001), JF-ASHS (Figure 3D; M=−174 voxels, CI=[−342, −5], *p*=0.044), and Infant-Trained-ASHS (Figure 3E; M=−463, CI=[−648, −279], *p*<0.001). Apart from Adult-Pretrained-ASHS (*r*=0.11, *p*=0.392), the ASHS models produced modest negative correlations (CE-ASHS: *r*=−0.39, *p*=0.013; JF-ASHS: *r*=−0.41, *p*=0.007; Infant-Trained-ASHS: *r*=−0.38, *p*=0.013), reflecting greater under-estimation for larger hippocampal volumes. Together, these bias data reveal notable differences between FreeSurfer and ASHS automated protocols.

## 4. Discussion

We compared four methods for segmenting the hippocampus from T1-weighted anatomical MRI scans in awake infants: average adult template, average infant template, FreeSurfer (Iglesias et al., 2015), and ASHS (Yushkevich et al., 2015). Each method depends on a library of manual segmentations from other brains (adults or infants), combined in different ways, to identify which voxels belong to the hippocampus in a new scan, without needing manual segmentation. In this paper, we conducted manual segmentation in order to evaluate the performance of each method but given our results that may not be necessary in future studies. Specifically, we quantified how well the segmentation from each method predicted the corresponding manual segmentation using Dice similarity. The manual segmentations also allowed us to create a gold-standard benchmark based on IRR between the two tracers, against which the Dice metrics for each method could be compared. FreeSurfer generated moderately accurate segmentations approaching IRR, but this came at the cost of a large bias to over-estimate hippocampal volume and to lose accuracy for greater volumes. ASHS outperformed the other methods in several ways, reliably predicting out-of-sample manual segmentations, exceeding IRR of manual segmentations, generalizing across tracers, and mostly avoiding strong bias. This indicates that automated algorithms developed for adults can nevertheless provide accurate and generalizable hippocampal segmentations in infants, especially when trained on infants.

The results illustrate the challenge of hippocampal segmentation in infants. Even though both tracers adhered to the same manual infant hippocampal segmentation protocol (Gousias et al., 2012), IRR was only moderate. The hippocampal head was particularly difficult to segment. The anterior extent of the hippocampus was ambiguous with low signal contrast, which led to inconsistency between the tracers. This is partly a consequence of collecting anatomical scans while infants are awake, and the use of an anatomical sequence not optimized for hippocampal segmentation (typical high-resolution turbo spin echo T2-weighted sequences present acoustic and specific absorption ratio risks to infants and are even more motion-sensitive).

The ASHS models trained on one tracer’s data excelled both within and across tracers. These models were trained in a leave-one-scan-out fashion to predict the manual segmentations from the tracer used for training and the tracer not used for training. In both tests and for both tracers, these ASHS models either exceeded or were similar to the manual IRR. Thus, ASHS trained on one tracer captures the idiosyncrasies of the data it was trained on without too much overfitting. Moreover, the ASHS model trained on data from both tracers produced segmentations that significantly outperformed IRR in predicting each tracer and most closely approximated what was optimally shared across tracers. We have publicly released this combined model and training data to help the community, as it will be most likely to succeed when applied to new infants and validated with new tracers.

ASHS was able to excel despite the noise in the infant scans because of the model’s ability to label “noncontroversial” voxels as hippocampus. This is built into the multi-atlas label fusion process in ASHS, which ensures that each training sample is used to ‘vote’ on how a voxel should be labeled, making sure that only high consensus voxels are labeled (Yushkevich et al., 2015). The ASHS models in the present study received 40+ manual segmentations of training data (whereas 20–30 are recommended), further helping ASHS form an accurate, reliable, and robust model for infant hippocampal segmentation.

Several limitations of the present study need to be acknowledged. Although ASHS proved successful, other methods have emerged in recent years that ostensibly outperform ASHS (Zhu et al., 2019). We are hopeful that our conclusions about generalization across tracers apply to those methods, but testing relative to IRR will be necessary. Another important consideration is that we evaluated methods with data from one, relatively impoverished MRI sequence not designed for hippocampal segmentation — short, quiet T1-weighted PETRA scans from awake infants. This was reasonable given our goal to assess which methods would work best for such data, as this is by far the anatomical sequence with the highest success rate in our awake infant studies (Ellis, et al., 2020). Future research will be needed to verify whether our findings apply to other sequences (e.g., T2-weighted MRI data from sleeping infants). That said, the ability of ASHS pre-trained with data from adults aged 55+ to accurately predict manual segmentations of the infant hippocampus suggests that differences in quality might not affect ASHS training severely.

Finally, given that the two tracers of this study segmented the whole infant hippocampus, the efficacy of ASHS for segmenting subfields with infant data was not assessed. This will be an important task for future research but awaits a clearer consensus in the field about anatomical landmarks for infant subfields. Such consensus in adults is what unleashed and enabled the rapid growth of structural and functional imaging of human hippocampus in recent years.

In conclusion, this study demonstrated the potential for automated algorithms to rival or replace expert tracers for hippocampal segmentation in infants. Although a nonlinearly aligned infant template and FreeSurfer modestly approached the IRR average, we present the first evidence that ASHS generates infant hippocampal segmentations that exceed IRR. Although other studies have shown the success of automated hippocampal segmentation in adults (Yushkevich et al., 2015), children (Schlichting et al., 2019), and infants (Guo et al., 2015; Zhu et al., 2019), this study is the first, to our knowledge, to show that automated methods for segmenting infant hippocampus can generalize across tracers and can work for scans collected in awake infants. As task-based, awake infant fMRI becomes more prevalent (Ellis et al., 2020; Ellis et al., 2021a, 2021b, 2021c), there will be increasing need for protocols that produce accurate and reliable segmentations of the infant hippocampus from noisy data. Automated methods make it possible to accelerate and improve the study of the developing hippocampus.

## Author Contributions

**Jared Fel:** Conceptualization, Methodology, Formal Analysis, Investigation, Writing – Original Draft Preparation, Writing - Review & Editing, Visualization. **Cameron Ellis:** Conceptualization, Methodology, Investigation, Writing - Review & Editing, Supervision. **Nicholas Turk-Browne:** Conceptualization, Writing - Review & Editing, Supervision, Funding Acquisition.

## Acknowledgements

This work would not be possible without the tremendous data collection and preprocessing efforts of T. Yates, L. Skalaban, and A. Bracher. We thank all the families who gave their time to participate in this study; K. Armstrong, L. Rait, J. Daniels, and the entire Yale Baby School team for recruitment, scheduling, and administration; and R. Lee, L. Nystrom, N. DePinto, and R. Watts for technical support. We are grateful for internal funding from the Department of Psychology and Princeton Neuroscience Institute at Princeton University and from the Department of Psychology and Faculty of Arts and Sciences at Yale University. N.B.T.-B. received further support from the Canadian Institute for Advanced Research and the James S. McDonnell Foundation (https://doi.org/10.37717/2020-1208).

## References

Aly, M. & Turk-Browne, N. B. (2016). Attention stabilizes representations in the human hippocampus. Cerebral Cortex, 26(2), 783–796.

Arnold, S. E. & Trojanowski, J. Q. (1996). Human fetal hippocampal development: I. cytoarchitecture, myeloarchitecture, and neuronal morphologic features. The Journal of Comparative Neurology, 367(2), 274–292.

Avants, B. B., Tustison, N. J., Song, G., Cook, P. A., Klein, A., & Gee, J. C. (2011). A reproducible evaluation of ants similarity metric performance in brain image registration. NeuroImage, 54(3), 2033–2044.

Bender, A. R., Keresztes, A., Bodammer, N. C., Shing, Y. L., Werkle-Bergner, M., Daugherty, A. M., Yu, Q., Kühn, S., Lindenberger, U., & Raz, N. (2018). Optimization and validation of automated hippocampal subfield segmentation across the lifespan. Human Brain Mapping, 39(2), 916–931.

Bland, J. M., & Altman, D. G. (2007). Agreement between methods of measurement with multiple observations per individual. Journal of Biopharmaceutical Statistics, 17(4), 571–582.

Boccardi, M., Bocchetta, M., Apostolova, L. G., Barnes, J., Bartzokis, G., Corbetta, G., DeCarli, C., deToledo-Morrell, L., Firbank, M., Ganzola, R., Gerritsen, L., Henneman, W., Killiany, R. J., Malykhin, N., Pasqualetti, P., Pruessner, J. C., Redolfi, A., Robitaille, N., Soininen, H., Tolomeo, D., … EADC-ADNI Working Group on the Harmonized Protocol for Manual Hippocampal Segmentation (2015). Delphi definition of the EADC-ADNI Harmonized Protocol for hippocampal segmentation on magnetic resonance. Alzheimer’s & Dementia: The Journal of the Alzheimer’s Association, 11(2), 126–138.

Brown, E. M., Pierce, M. E., Clark, D. C., Fischl, B. R., Iglesias, J. E., Milberg, W. P., McGlinchey, R. E., & Salat, D. H. (2020). Test-retest reliability of FreeSurfer automated hippocampal subfield segmentation within and across scanners. NeuroImage, 210, 116563.

Carmo, D., Silva, B., Yasuda, C., Rittner, L., & Lotufo, R. (2019). Extended 2D volumetric consensus hippocampus segmentation. ArXiv, abs/1902.04487.

Cohen N. J. & Eichenbaum H. (1993). Memory, Amnesia, and the Hippocampal System. Cambridge, MA, MIT Press.

Colon-Perez, L. M., Triplett, W., Bohsali, A., Corti, M., Nguyen, P. T., Patten, C., Mareci, T. H., & Price, C. C. (2016). A majority rule approach for region-of-interest-guided streamline fiber tractography. Brain Imaging and Behavior, 10(4), 1137–1147.

Corkin, S. (2013). Permanent Present Tense: The Unforgettable Life of the Amnesic Patient, HM. Basic Books (AZ).

Dalton, M. A., Zeidman, P., Barry, D. N., Williams, E., & Maguire, E. A. (2017). Segmenting subregions of the human hippocampus on structural magnetic resonance image scans: An illustrated tutorial. Brain and Neuroscience Advances, 1, 2398212817701448.

DeMaster, D., Pathman, T., Lee, J. K., & Ghetti, S. (2014). Structural development of the hippocampus and episodic memory: developmental differences along the anterior/posterior axis. Cerebral Cortex, 24(11), 3036–3045.

Denisova K. (2019). Age attenuates noise and increases symmetry of head movements during sleep resting-state fMRI in healthy neonates, infants, and toddlers. Infant Behavior & Development, 57, 101317.

Deoni, S., Bruchhage, M., Beauchemin, J., Volpe, A., D’Sa, V., Huentelman, M., & Williams, S. (2021). Accessible pediatric neuroimaging using a low field strength MRI scanner. NeuroImage, 238, 118273.

Dice, L. R. (1945). Measures of the amount of ecologic association between species. Ecology, 26(3), 297–302.

Efron, B., & Tibshirani, R. (1986). Bootstrap methods for standard errors, confidence intervals, and other measures of statistical accuracy. Statistical Science, 1(1), 54–75.

Ellis, C. T., Skalaban, L. J., Yates, T. S., Bejjanki, V. R., Córdova, N. I., & Turk-Browne, N. B. (2021a). Evidence of hippocampal learning in human infants. Current Biology, 31, 1–7.

Ellis, C. T., Yates, T. S., Skalaban, L. J., Bejjanki, V. R., Arcaro, M. J., & Turk-Browne, N. B. (2021b). Retinotopic organization of visual cortex in human infants. Neuron, 109, 1–11.

Ellis, C. T., Skalaban, L. J., Yates, T. S., & Turk-Browne. N. B. (2021c). Attention recruits frontal cortex in human infants. Proceedings of the National Academia of Sciences, 118(12), e2021474118.

Ellis, C. T., Skalaban, L. J., Yates, T. S., Bejjanki, V. R., Cordova, N. I., & Turk-Browne, N.B. (2020). Re-imagining fMRI for awake behaving infants. Nature Communications, 11, 4523.

Fonov, V., Evans, A. C., Botteron, K., Almli, C. R., McKinstry, R. C., Collins, D. L., & Brain Development Cooperative Group (2011). Unbiased average age-appropriate atlases for pediatric studies. NeuroImage, 54(1), 313–327.

Frisoni, G. B., Jack, C. R., Jr, Bocchetta, M., Bauer, C., Frederiksen, K. S., Liu, Y., Preboske, G., Swihart, T., Blair, M., Cavedo, E., Grothe, M. J., Lanfredi, M., Martinez, O., Nishikawa, M., Portegies, M., Stoub, T., Ward, C., Apostolova, L. G., Ganzola, R., Wolf, D., … EADC-ADNI Working Group on The Harmonized Protocol for Manual Hippocampal Volumetry and for the Alzheimer’s Disease Neuroimaging Initiative (2015). The EADC-ADNI Harmonized Protocol for manual hippocampal segmentation on magnetic resonance: evidence of validity. Alzheimer’s & Dementia, 11(2), 111–125.

Gogtay, N., Nugent, T. F., 3rd, Herman, D. H., Ordonez, A., Greenstein, D., Hayashi, K. M., Clasen, L., Toga, A. W., Giedd, J. N., Rapoport, J. L., & Thompson, P. M. (2006). Dynamic mapping of normal human hippocampal development. Hippocampus, 16(8), 664–672.

Gómez, R. L., & Edgin, J. O. (2016). The extended trajectory of hippocampal development: Implications for early memory development and disorder. Developmental Cognitive Neuroscience, 18, 57–69.

Gousias, I. S., Edwards, A. D., Rutherford, M. A., Counsell, S. J., Hajnal, J. V., Rueckert, D., & Hammers, A. (2012). Magnetic resonance imaging of the newborn brain: manual segmentation of labelled atlases in term-born and preterm infants. NeuroImage, 62(3), 1499–1509.

Guo, T., Winterburn, J. L., Pipitone, J., Duerden, E. G., Park, M. T., Chau, V., Poskitt, K. J., Grunau, R. E., Synnes, A., Miller, S. P., & Mallar Chakravarty, M. (2015). Automatic segmentation of the hippocampus for preterm neonates from early-in-life to term-equivalent age. NeuroImage: Clinical, 9, 176–193.

Guo, Y., Wu, G., Commander, L. A., Szary, S., Jewells, V., Lin, W., & Shent, D. (2014). Segmenting hippocampus from infant brains by sparse patch matching with deep-learned features. International Conference on Medical Image Computing and Computer-Assisted Intervention, 17(Pt 2), 308–315.

Hashempour, N., Tuulari, J. J., Merisaari, H., Lidauer, K., Luukkonen, I., Saunavaara, J., Parkkola, R., Lähdesmäki, T., Lehtola, S. J., Keskinen, M., Lewis, J. D., Scheinin, N. M., Karlsson, L., & Karlsson, H. (2019). A novel approach for manual segmentation of the amygdala and hippocampus in neonate MRI. Frontiers in Neuroscience, 13, 1025.

Iglesias, J. E., Augustinack, J. C., Nguyen, K., Player, C. M., Player, A., Wright, M., Roy, N., Frosch, M. P., McKee, A. C., Wald, L. L., Fischl, B., Van Leemput, K., & Alzheimer’s Disease Neuroimaging Initiative (2015). A computational atlas of the hippocampal formation using ex vivo, ultra-high resolution MRI: Application to adaptive segmentation of in vivo MRI. NeuroImage, 115, 117–137.

Insausti, R., & Amaral, D. G. (2004). Hippocampal formation. In The Human Nervous System: Second Edition, 871–914.

Insausti, R., Cebada-Sánchez, S., & Marcos, P. (2010). Postnatal development of the human hippocampal formation. Advances in Anatomy, Embryology, and Cell Biology, 206, 1–86.

Keresztes, A., Ngo, C. T., Lindenberger, U., Werkle-Bergner, M., & Newcombe, N. S. (2018). Hippocampal maturation drives memory from generalization to specificity. Trends in Cognitive Sciences, 22(8), 676–686.

Khlif, M. S., Egorova, N., Werden, E., Redolfi, A., Boccardi, M., DeCarli, C. S., Fletcher, E., Singh, B., Li, Q., Bird, L., & Brodtmann, A. (2019). A comparison of automated segmentation and manual tracing in estimating hippocampal volume in ischemic stroke and healthy control participants. NeuroImage: Clinical, 21, 101581.

Li, G., Chen, M. H., Li, G., Wu, D., Sun, Q., Shen, D., & Wang, L. (2019). A preliminary volumetric MRI study of amygdala and hippocampal subfields in autism during infancy. Proceedings. IEEE International Symposium on Biomedical Imaging, 2019, 1052–1056.

Makris, N., Goldstein, J. M., Kennedy, D., Hodge, S. M., Caviness, V. S., Faraone, S. V., Tsuang, M. T., & Seidman, L. J. (2006). Decreased volume of left and total anterior insular lobule in schizophrenia. Schizophrenia Research, 83(2-3), 155–171.

Mange, J. (2019). Effect of training data order for machine learning. 2019 International Conference on Computational Science and Computational Intelligence (CSCI), 2019, 406–407.

Mazziotta, J., Toga, A., Evans, A., Fox, P., Lancaster, J., Zilles, K., Woods, R., Paus, T., Simpson, G., Pike, B., Holmes, C., Collins, L., Thompson, P., MacDonald, D., Iacoboni, M., Schormann, T., Amunts, K., Palomero-Gallagher, N., Geyer, S., Parsons, L., … Mazoyer, B. (2001). A probabilistic atlas and reference system for the human brain: International Consortium for Brain Mapping (ICBM). Philosophical Transactions of the Royal Society of London. Series B, Biological Sciences, 356(1412), 1293–1322.

Morey, R. A., Petty, C. M., Xu, Y., Hayes, J. P., Wagner, H. R., 2nd, Lewis, D. V., LaBar, K. S., Styner, M., & McCarthy, G. (2009). A comparison of automated segmentation and manual tracing for quantifying hippocampal and amygdala volumes. NeuroImage, 45(3), 855–866.

Nadel, L., & Hardt, O. (2011). Update on memory systems and processes. Neuropsychopharmacology, 36(1), 251–273.

Pineda, R. G., Neil, J., Dierker, D., Smyser, C. D., Wallendorf, M., Kidokoro, H., Reynolds, L. C., Walker, S., Rogers, C., Mathur, A. M., Van Essen, D. C., & Inder, T. (2014). Alterations in brain structure and neurodevelopmental outcome in preterm infants hospitalized in different neonatal intensive care unit environments. Journal of Pediatrics, 164(1), 52–60.e2.

Rakic, P., & Nowakowski, R. S. (1981). The time of origin of neurons in the hippocampal region of the rhesus monkey. Journal of Comparative Neurology, 196(1), 99–128.

Rodionov, R., Chupin, M., Williams, E., Hammers, A., Kesavadas, C., & Lemieux, L. (2009). Evaluation of atlas-based segmentation of hippocampi in healthy humans. Magnetic Resonance Imaging, 27(8), 1104–1109.

Schapiro, A. C., Turk-Browne, N. B., Botvinick, M. M., & Norman, K. A. (2017). Complementary learning systems within the hippocampus: a neural network modelling approach to reconciling episodic memory with statistical learning. Philosophical Transactions of the Royal Society of London. Series B, Biological Sciences, 372(1711), 20160049.

Schlichting, M. L., Guarino, K. F., Schapiro, A. C., Turk-Browne, N. B., & Preston, A. R. (2017). Hippocampal structure predicts statistical learning and associative inference abilities during development. Journal of Cognitive Neuroscience, 29(1), 37–51.

Schlichting, M. L., Mack, M. L., Guarino, K. F., & Preston, A. R. (2019). Performance of semi-automated hippocampal subfield segmentation methods across ages in a pediatric sample. NeuroImage, 191, 49–67.

Schmidt, M. F., Storrs, J. M., Freeman, K. B., Jack, C. R., Jr, Turner, S. T., Griswold, M. E., & Mosley, T. H., Jr (2018). A comparison of manual tracing and FreeSurfer for estimating hippocampal volume over the adult lifespan. Human Brain Mapping, 39(6), 2500–2513.

Schoemaker, D., Buss, C., Head, K., Sandman, C. A., Davis, E. P., Chakravarty, M. M., Gauthier, S., & Pruessner, J. C. (2016). Hippocampus and amygdala volumes from magnetic resonance images in children: Assessing accuracy of FreeSurfer and FSL against manual segmentation. NeuroImage, 129, 1–14.

Shi, F., Yap, P. T., Wu, G., Jia, H., Gilmore, J. H., Lin, W., & Shen, D. (2011). Infant brain atlases from neonates to 1- and 2-year-olds. PloS One, 6(4), e18746.

Squire L. R. (1992). Memory and the hippocampus: a synthesis from findings with rats, monkeys, and humans. Psychological Review, 99(2), 195–231.

Tulving, E., & Markowitsch, H. J. (1998). Episodic and declarative memory: role of the hippocampus. Hippocampus, 8(3), 198–204.

Uematsu, A., Matsui, M., Tanaka, C., Takahashi, T., Noguchi, K., Suzuki, M., & Nishijo, H. (2012). Developmental trajectories of amygdala and hippocampus from infancy to early adulthood in healthy individuals. PloS One, 7(10), e46970.

Xie, L., Wisse, L., Pluta, J., de Flores, R., Piskin, V., Manjón, J. V., Wang, H., Das, S. R., Ding, S. L., Wolk, D. A., Yushkevich, P. A., & Alzheimer’s Disease Neuroimaging Initiative (2019). Automated segmentation of medial temporal lobe subregions on in vivo T1-weighted MRI in early stages of Alzheimer’s disease. Human Brain Mapping, 40(12), 3431–3451.

Xie, L., Wisse, L., Das, S. R., Wang, H., Wolk, D. A., Manjón, J. V., & Yushkevich, P. A. (2016). Accounting for the confound of meninges in segmenting entorhinal and perirhinal cortices in T1-Weighted MRI. International Conference on Medical Image Computing and Computer-Assisted Intervention, 9901, 564–571.

Xue, H., Srinivasan, L., Jiang, S., Rutherford, M., Edwards, A. D., Rueckert, D., & Hajnal, J. V. (2007). Automatic segmentation and reconstruction of the cortex from neonatal MRI. NeuroImage, 38(3), 461–477.

Yushkevich, P. A., Pluta, J. B., Wang, H., Xie, L., Ding, S. L., Gertje, E. C., Mancuso, L., Kliot, D., Das, S. R., & Wolk, D. A. (2015). Automated volumetry and regional thickness analysis of hippocampal subfields and medial temporal cortical structures in mild cognitive impairment. Human Brain Mapping, 36(1), 258–287.

Zhu, H., Shi, F., Wang, L., Hung, S. C., Chen, M. H., Wang, S., Lin, W., & Shen, D. (2019). Dilated dense U-Net for infant hippocampus subfield segmentation. Frontiers in Neuroinformatics, 13, 30.

