## Supplementary Information for "Automated and manual segmentation of the hippocampus in human infants"

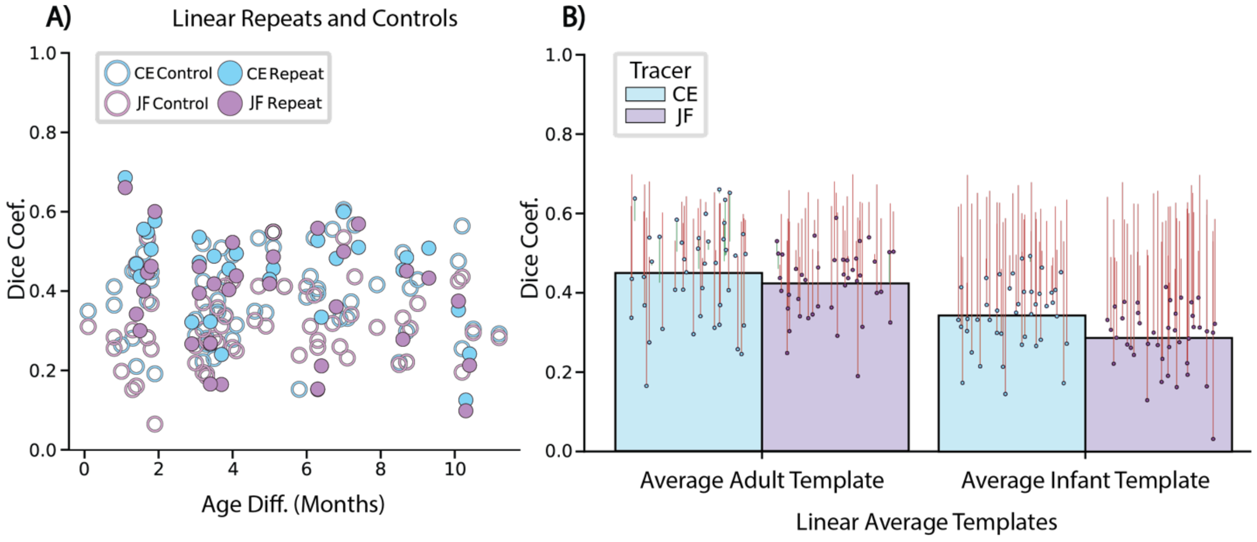


**Figure S1. Linear repeat, control, and average template data.** (A) Segmentations aligned across sessions using linear transformation methods described previously (Ellis et al., 2020): The y-axis represents the Dice between the paired sessions. The x-axis represents the difference in age between sessions being compared. The average age match error between the repeat and control pairs was 0.31 months (SD=0.38; range=0.00–1.40). Repeat Dice comparisons were significantly greater than corresponding control Dice comparisons (CE: repeat Dice=0.45, control Dice=0.39, M=0.06, CI=[0.020, 0.092], $p$=0.003; JF: repeat Dice=0.39, control Dice=0.32, M=0.07, CI=[0.038, 0.110], $p$<0.001). The repeat Dice values of both tracers were also negatively correlated with the age difference (CE repeat: $r$=-0.39, $p$=0.003; CE control: $r$=0.10, $p$=0.443; JF repeat: $r$=-0.27, $p$=0.042; JF control: $r$=0.13, $p$=0.306) — as the age difference of the two repeat sessions decreased, accuracy increased. (B) Linear average templates predicting manual hippocampal segmentations: Each colored dot is one participant scan. Each colored line extending from the dots displays how much improvement (green) or deterioration (red) between the similarity to template and the IRR for that session. The average Dice between the linear average adult template and the linearly aligned manual infant segmentations was low for both tracers (Table S3; CE: Dice=0.45, SD=0.12, range=0.16– 0.66; JF: Dice=0.42, SD=0.09, range=0.19–0.29) and significantly underperformed the manual IRR average of 0.57 (CE: M=-0.12, CI=[-0.168, -0.079], p<0.001; JF: M=-0.15, CI=[-0.184, -0.115], p<0.001). Unexpectedly, the linear average infant template had a lower similarity for both tracers than the average adult template (Table S3; CE: Dice=0.34, SD=0.09, range=0.15– 0.49; JF: Dice=0.29, SD=0.08, range=0.03–0.42) and was significantly worse than the manual IRR (CE: M=-0.23, CI=[-0.263, -0.194], p<0.001; JF: M=-0.28, CI=[-0.315, -0.256], p<0.001). The success of the average adult template over the average infant template suggests that linearly aligning infant data to a standard space is hurting precision.

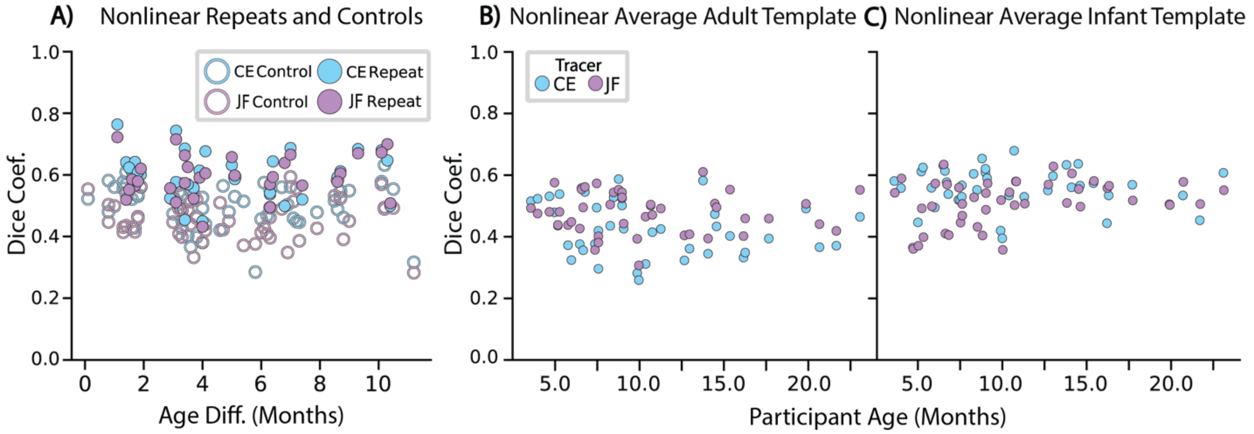


**Figure S2. Nonlinear repeat, control, and average template correlations.** (A) Nonlinear repeat and control hippocampal Dices compared with the age difference of the two scans. The y-axis represents the Dice between the paired sessions. The x-axis represents the difference in age between sessions being compared. The average age match error between the repeat and control pairs was 0.31 months (SD=0.38; range=0.00–1.40). Both CE and JF's hippocampal repeat means were significantly higher than respective controls (CE: repeat Dice=0.60, control Dice=0.51, M=0.10, CI=[0.077, 0.116], $p$<0.001; JF: repeat Dice=0.60, control Dice=0.47, M=0.13, CI=[0.105, 0.148], $p$<0.001). There were no reliable correlations between Dice and age difference (CE repeat: $r$=-0.10, $p$=0.428; CE control: $r$=-0.15, $p$=0.285; JF repeat: $r$=0.16, $p$=0.248; JF control: $r$=-0.08, $p$=0.621). Repeat sessions were more similar when using nonlinear alignment methods than linear alignment methods for both tracers (CE: M=0.16, CI=[0.123, 0.189], $p$<0.001; JF: M=0.20, CI=[0.169, 0.241], $p$<0.001). (B) No reliable correlations with age were observed for the accuracy of the average adult template in approximating the nonlinearly transformed manual segmentations (CE: $r$=-0.23, $p$=0.063; JF: $r$=-0.03, $p$=0.797). (C) A positive correlation between age and accuracy of the nonlinear infant template was found only for JF (CE: $r$=-0.03, $p$=0.811; JF: $r$=0.32, $p$=0.008).

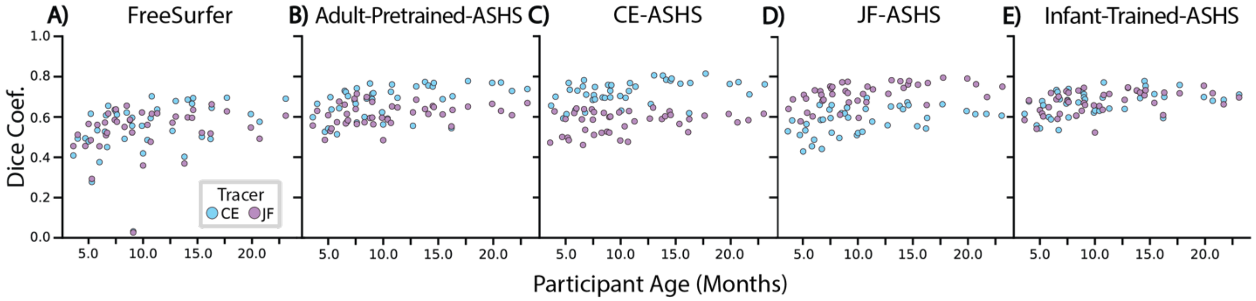


**Figure S3. Age correlations with the accuracy of FreeSurfer and ASHS models.** When Dice values were plotted against infant age, there were significant positive correlations with the predictions of both tracers for every model: (A) FreeSurfer (CE: $r$=0.36, $p$=0.002; JF: $r$=0.22, $p$=0.032), (B) Adult-Pretrained-ASHS (CE: $r$=0.47, $p$=0.001; JF: $r$=0.30, $p$=0.019), (C) CE-ASHS (CE: $r$=0.44, $p$=0.002; JF: $r$=0.26, $p$=0.040), (D) JF-ASHS (CE: $r$=0.42, $p$<0.001; JF: $r$=0.50, $p$<0.001), and Infant-Trained-ASHS (CE: $r$=0.40, $p$=0.001; JF: $r$=0.42, $p$<0.001).

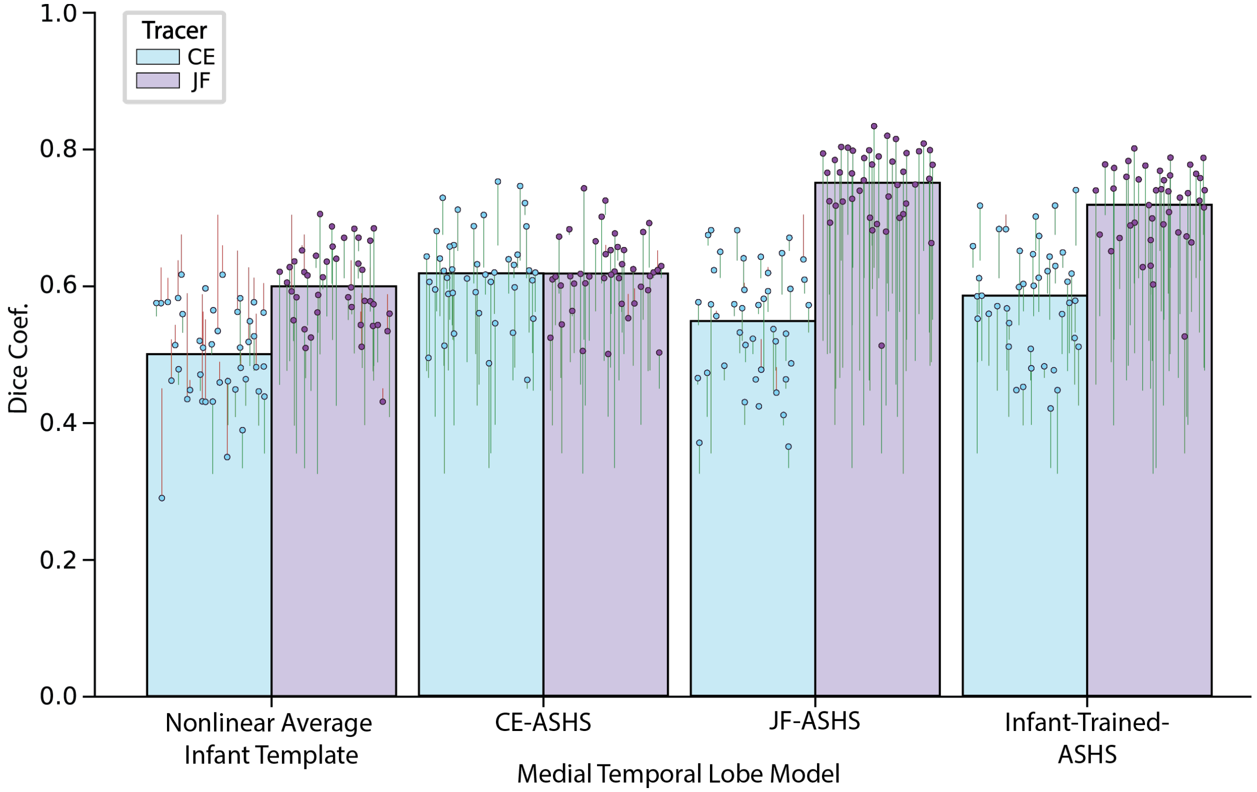


**Figure S4. Medial temporal lobe (MTL) cortex results.** Each colored dot is one participant scan. Each colored line extending from the dots depicts the improvement (green) or deterioration (red) between the similarity to template/ASHS model and the between-tracer MTL cortex IRR of 0.52. A nonlinear infant MTL cortex template (CE: Dice=0.50, SD=0.07, range=0.29–0.61; JF: Dice=0.59, SD=0.06, range=0.43–0.70) predicted CE's data similarly to IRR (M=-0.02, CI=[-0.042, 0.000], $p$=0.055) and significantly exceeded IRR in predicting JF's data (M=0.08, CI=[0.049, 0.105], $p$<0.001). The template's predictions correlated with age for each tracer (CE: $r$=0.34, $p$=0.017; JF: $r$=0.36, $p$=0.018). Both the CE-ASHS and JF-ASHS models predicted held-out manual segmentations from the same tracer (CE-ASHS predicting CE: Dice=0.61, SD=0.07, range=0.46–0.75; JF-ASHS predicting JF: Dice=0.74, SD=0.06, range=0.51–0.83) with an accuracy that significantly exceeded IRR (CE: M=0.10, CI=[0.080, 0.113], $p$<0.001; JF: M=0.23, CI=[0.202, 0.253], $p$<0.001). Both models predicted the tracer they were not trained on (JF-ASHS predicting CE: Dice=0.54, SD=0.09, range=0.36–0.68; CE-ASHS predicting JF: Dice=0.61, SD=0.06, range=0.50–0.74) better than the other tracer, i.e., significantly surpassing IRR (CE: M=0.03, CI=[0.014, 0.041], $p$<0.001; JF: M=0.10, CI=[0.073, 0.118], $p$<0.001). When Dice values were plotted against infant age, the accuracy of the CE-trained model was significantly positively correlated for both tracers, (CE: $r$=0.43, $p$<0.001; JF: $r$=0.55, $p$<0.001), while the performance of the JF-ASHS model did not generate significant positive correlations in its approximations of either tracer's MTL cortex data (CE: $r$=0.28, $p$=0.076; JF: $r$=0.03, $p$=0.773). Moreover, the Infant-Trained-ASHS model predicted both tracers (CE: Dice=0.58, SD=0.08, range=0.42–0.73; JF: Dice=0.71; SD=0.06, range=0.52–0.79) better than IRR (CE: M=0.06, CI=[0.051, 0.077], $p$<0.001; JF: M=0.20, CI=[0.173, 0.219], $p$<0.001) and in a way that correlated weakly with age (CE: $r$=0.33, $p$=0.033; JF: $r$=0.24, $p$=0.083). Overall, this pattern of results closely mirrored the hippocampus.

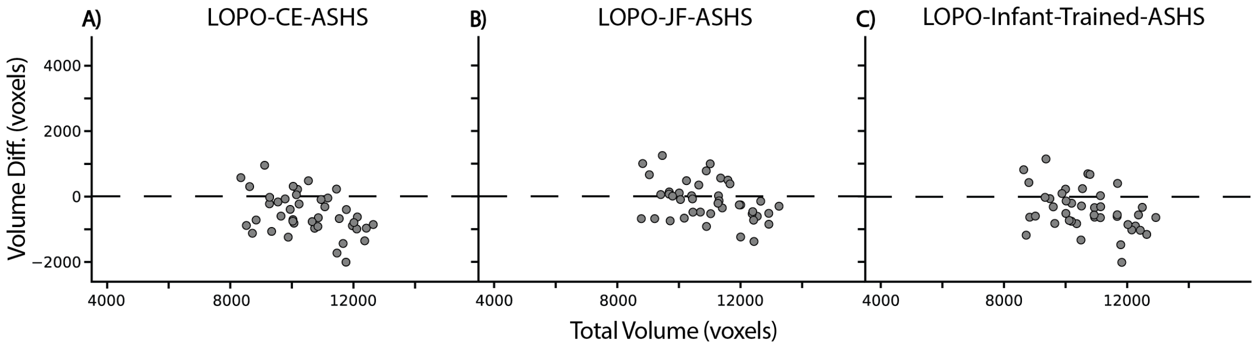


**Figure S5. Leave-one-participant-out (LOPO) ASHS hippocampal bias plots.** Consistent with the leave-one-scan-out ASHS models, the (A) LOPO-CE-ASHS model (M=-546 voxels, CI=[-736, -356], p<0.001) and (C) LOPO-Infant-Trained-ASHS model (M=-420, CI=[-616, -230], p<0.001) under-estimated hippocampal volume, whereas the (B) LOPO-JF-ASHS model (M=-149 voxels, CI=[-327, 29], p=0.103) did not. All three LOPO ASHS models showed a reliable bias correlation (LOPO-CE-ASHS: *r*=-0.42, p=0.005; LOPO-JF-ASHS: *r*=-0.40, p=0.010; LOPO-Infant-Trained-ASHS: *r*=-0.40, p=0.009), with greater under-estimation for participants with larger hippocampal volumes. These moderate correlations further reinforce the contrast in bias between FreeSurfer and ASHS.

| ASHS Model | CE HPC | | | JF HPC | | | CE MTL | | | JF MTL | | |
|---|---|---|---|---|---|---|---|---|---|---|---|---|
| | **Dice** | **SD** | **Range** | **Dice** | **SD** | **Range** | **Dice** | **SD** | **Range** | **Dice** | **SD** | **Range** |
| CE-ASHS | 0.73 | 0.06 | 0.59–0.82 | 0.58 | 0.05 | 0.46–0.65 | 0.61 | 0.07 | 0.46–0.75 | 0.61 | 0.06 | 0.50–0.74 |
| LOPO-CE-ASHS | 0.72 | 0.06 | 0.60–0.81 | 0.58 | 0.05 | 0.48–0.65 | 0.61 | 0.07 | 0.46–0.74 | 0.61 | 0.05 | 0.49–0.73 |
| JF-ASHS | 0.59 | 0.07 | 0.43–0.70 | 0.72 | 0.06 | 0.58–0.80 | 0.54 | 0.09 | 0.36–0.68 | 0.74 | 0.06 | 0.51–0.83 |
| LOPO-JF-ASHS | 0.59 | 0.07 | 0.43–0.69 | 0.71 | 0.06 | 0.58–0.80 | 0.54 | 0.09 | 0.36–0.68 | 0.74 | 0.06 | 0.51–0.82 |
| Infant-Trained-ASHS | 0.67 | 0.06 | 0.54–0.78 | 0.67 | 0.06 | 0.52–0.76 | 0.58 | 0.08 | 0.42–0.73 | 0.71 | 0.06 | 0.52–0.79 |
| LOPO-Infant-Trained-ASHS | 0.67 | 0.06 | 0.47–0.78 | 0.67 | 0.06 | 0.55–0.77 | 0.58 | 0.08 | 0.44–0.73 | 0.71 | 0.05 | 0.55–0.79 |

**Table S1. Leave-one-participant-out (LOPO) ASHS.** LOPO-CE-ASHS, LOPO-JF-ASHS, and LOPO-Infant-Trained-ASHS Dice values were generated by leaving out all sessions from one participant during training, rather than just the individual session being tested. In all three models, for every ROI, and from each tracer, the performance of the LOPO-ASHS model was either slightly lower or the same, on average, compared to its leave-one-scan-out ASHS model counterpart. Instances of a reduction in Dice were expected because less data were used for training. Overall, this analysis shows that the protocols can survive stricter tests of independence.

| ID | Age | Sex | Location | Merged | State |
|---|---|---|---|---|---|
| s6057_1_1 | 3.6 | M | BIC | No | A |
| s6607_1_1 | 4 | M | MRRC | No | A |
| s7067_1_1 | 4.7 | F | BIC | No | A |
| s6687_1_1 | 5 | F | MRRC | No | A |
| s7017_1_1 | 5.2 | F | BIC | No | A |
| s0687_1_1 | 5.3 | F | MRRC | No | A |
| s8687_1_2 | 5.8 | F | MRRC | No | A |
| s2687_1_2 | 6 | M | MRRC | Yes | S |
| s2037_1_2 | 6.5 | M | BIC | No | A |
| s7017_1_2 | 6.6 | F | BIC | No | S |
| s4607_1_2 | 6.7 | F | MRRC | No | A |
| s1057_1_1 | 6.8 | F | BIC | No | A |
| s6607_1_2 | 7.4 | M | BIC | No | A |
| s3607_1_1 | 7.5 | F | MRRC | No | S |
| s0607_1_1 | 7.6 | M | MRRC | No | A |
| s6057_1_2 | 7.6 | M | BIC | No | A |
| s7017_1_4 | 8.3 | F | BIC | No | A |
| s8607_1_1 | 8.5 | F | MRRC | No | A |
| s4107_1_1 | 8.8 | M | P.ton | Yes | S |
| s0687_1_2 | 9 | F | MRRC | No | S |
| s0057_1_3 | 9 | F | BIC | No | A |
| s0307_1_2 | 9.1 | M | P.ton | Yes | S |
| s2687_1_3 | 9.9 | M | BIC | No | A |
| s1017_1_1 | 10 | F | BIC | No | A |
| s3607_1_2 | 10.4 | F | BIC | No | A |
| s1607_1_2 | 10.7 | M | BIC | No | A |
| s6607_1_3 | 10.8 | M | BIC | No | A |
| s6687_1_3 | 11.3 | F | BIC | No | A |
| s0607_1_2 | 12.7 | M | BIC | No | A |
| s4607_1_4 | 13 | F | BIC | Yes | A |
| s8187_1_4 | 13.8 | F | P.ton | Yes | S |
| s4607_1_5 | 14.1 | F | BIC | No | A |
| s8687_1_4 | 14.5 | F | BIC | Yes | A |
| s0607_1_3 | 14.6 | M | BIC | No | A |
| s6687_1_4 | 15.4 | F | BIC | Yes | A |
| s0607_1_4 | 16.2 | M | BIC | No | A |
| s2687_1_4 | 16.3 | M | BIC | No | A |
| s0607_1_5 | 17.7 | M | BIC | No | A |
| s2307_1_1 | 19.9 | M | P.ton | No | S |
| s1187_1_1 | 20.7 | F | P.ton | Yes | A |
| s2307_1_2 | 21.7 | M | P.ton | Yes | A |
| s8187_1_8 | 23.1 | F | P.ton | Yes | A |

**Table S2. Demographic information.** ‘ID’ is a unique infant identifier (i.e., sXXXX_Y_Z): the first four numbers (XXXX) indicate the family, the fifth number (Y) the infant number within the family, and the sixth number (Z) the session number for that infant. ‘Age’ is measured in months. ‘Sex’ is male or female. Location is 1.) Scully Center for the Neuroscience of Mind and Behavior at Princeton University (P.ton), 2.) Brain Imaging Center at Yale University (BIC), 3.) Magnetic Resonance Research Center at Yale University (MRRC). ‘Merged’ shows whether the scan was formed by aligning and averaging two scans together. ‘State’ indicates whether the infant was awake (A) or asleep (S) during the anatomical scan.

| ID | Age | IRR | Lin. Average Infant Template | | Lin. Average Adult Template | | Nonlin. Average Infant Template | | Nonlin. Average Adult Template | | FreeSurfer | |
|---|---|---|---|---|---|---|---|---|---|---|---|---|
| | | | CE | JF | CE | JF | CE | JF | CE | JF | CE | JF |
| s6057_1_1 | 3.6 | 0.50 | 0.49 | 0.38 | **0.58** | 0.46 | **0.58** | **0.54** | 0.51 | 0.49 | 0.41 | 0.46 |
| s6607_1_1 | 4.0 | 0.59 | 0.40 | 0.38 | 0.53 | 0.44 | 0.55 | 0.58 | 0.52 | 0.47 | 0.50 | 0.52 |
| s7067_1_1 | 4.7 | 0.46 | 0.30 | 0.31 | **0.54** | **0.50** | 0.36 | 0.36 | **0.53** | **0.48** | **0.50** | 0.46 |
| s6687_1_1 | 5.0 | 0.49 | 0.17 | 0.31 | 0.26 | 0.44 | 0.44 | 0.37 | 0.47 | 0.57 | 0.48 | **0.57** |
| s7017_1_1 | 5.2 | 0.48 | **0.49** | 0.29 | **0.58** | 0.45 | **0.61** | **0.49** | 0.43 | 0.43 | **0.62** | **0.49** |
| s0687_1_1 | 5.3 | 0.49 | 0.44 | 0.39 | 0.41 | 0.43 | **0.62** | 0.39 | **0.53** | 0.48 | 0.28 | 0.29 |
| s8687_1_2 | 5.8 | 0.57 | 0.30 | 0.27 | 0.41 | 0.38 | 0.55 | 0.49 | 0.37 | 0.43 | 0.54 | 0.57 |
| s2687_1_2 | 6.0 | 0.59 | 0.22 | 0.22 | 0.29 | 0.36 | 0.49 | 0.57 | 0.32 | 0.44 | 0.38 | 0.46 |
| s2037_1_2 | 6.5 | 0.64 | 0.37 | 0.33 | 0.45 | 0.48 | 0.62 | 0.63 | 0.37 | 0.42 | 0.53 | 0.58 |
| s7017_1_2 | 6.6 | 0.51 | 0.34 | 0.13 | **0.53** | 0.34 | **0.61** | 0.41 | **0.55** | **0.55** | **0.61** | **0.51** |
| s4607_1_2 | 6.7 | 0.49 | 0.38 | 0.37 | 0.27 | 0.34 | **0.57** | **0.56** | 0.35 | **0.49** | 0.45 | **0.53** |
| s1057_1_1 | 6.8 | 0.58 | 0.21 | 0.23 | 0.24 | 0.30 | 0.53 | 0.40 | 0.54 | 0.56 | **0.66** | **0.62** |
| s6607_1_2 | 7.4 | 0.63 | 0.25 | 0.24 | 0.37 | 0.31 | 0.51 | 0.55 | 0.37 | 0.35 | **0.64** | 0.59 |
| s3607_1_1 | 7.5 | 0.59 | 0.27 | 0.32 | 0.32 | 0.40 | 0.51 | 0.44 | 0.49 | 0.57 | **0.63** | **0.64** |
| s0607_1_1 | 7.6 | 0.68 | 0.40 | 0.34 | 0.54 | 0.48 | 0.56 | 0.46 | 0.42 | 0.40 | 0.62 | 0.58 |
| s6057_1_2 | 7.6 | 0.51 | 0.46 | 0.39 | 0.47 | 0.46 | **0.52** | 0.50 | 0.29 | 0.38 | nan | nan |
| s7017_1_4 | 8.3 | 0.50 | 0.41 | 0.27 | 0.49 | **0.50** | **0.60** | **0.52** | 0.43 | **0.50** | **0.50** | **0.56** |
| s8607_1_1 | 8.5 | 0.65 | 0.33 | 0.34 | 0.31 | 0.34 | 0.55 | 0.43 | 0.52 | 0.54 | 0.63 | **0.66** |
| s4107_1_1 | 8.8 | 0.53 | 0.17 | 0.03 | 0.41 | 0.25 | **0.65** | 0.40 | **0.58** | **0.55** | **0.59** | **0.59** |
| s0687_1_2 | 9.0 | 0.63 | 0.36 | 0.19 | 0.53 | 0.44 | 0.60 | 0.54 | 0.50 | 0.54 | 0.56 | 0.53 |
| s0057_1_3 | 9.0 | 0.56 | 0.35 | 0.30 | 0.47 | 0.42 | **0.61** | 0.48 | 0.52 | 0.52 | **0.60** | 0.53 |
| s0307_1_2 | 9.1 | 0.59 | 0.15 | 0.16 | 0.16 | 0.19 | 0.58 | 0.57 | 0.42 | 0.44 | 0.03 | 0.03 |
| s2687_1_3 | 9.9 | 0.60 | 0.37 | 0.31 | 0.44 | 0.50 | 0.42 | 0.51 | 0.28 | 0.39 | 0.56 | **0.63** |
| s1017_1_1 | 10.0 | 0.43 | 0.30 | 0.26 | **0.54** | **0.50** | 0.39 | 0.35 | 0.26 | 0.30 | 0.42 | 0.36 |
| s3607_1_2 | 10.4 | 0.50 | 0.35 | 0.26 | **0.51** | 0.48 | **0.53** | **0.54** | 0.31 | 0.46 | 0.49 | **0.57** |
| s1607_1_2 | 10.7 | 0.53 | 0.40 | 0.18 | **0.65** | 0.49 | **0.67** | 0.50 | 0.50 | 0.50 | **0.60** | 0.48 |
| s6607_1_3 | 10.8 | 0.66 | 0.33 | 0.27 | 0.41 | 0.39 | 0.57 | 0.57 | 0.41 | 0.47 | **0.71** | 0.62 |
| s6687_1_3 | 11.3 | 0.60 | 0.29 | 0.16 | 0.55 | 0.40 | 0.52 | 0.50 | 0.42 | 0.49 | **0.64** | **0.62** |
| s0607_1_2 | 12.7 | 0.58 | 0.45 | 0.42 | 0.50 | 0.54 | 0.55 | 0.56 | 0.32 | 0.40 | 0.54 | **0.58** |
| s4607_1_4 | 13.0 | 0.67 | 0.37 | 0.38 | 0.37 | 0.40 | 0.59 | 0.62 | 0.36 | 0.40 | **0.69** | 0.61 |
| s8187_1_4 | 13.8 | 0.65 | 0.45 | 0.30 | **0.66** | 0.59 | 0.63 | 0.50 | 0.58 | 0.60 | 0.41 | 0.37 |
| s4607_1_5 | 14.1 | 0.62 | 0.32 | 0.35 | 0.32 | 0.32 | 0.55 | 0.60 | 0.34 | 0.39 | **0.69** | 0.60 |
| s8687_1_4 | 14.5 | 0.70 | 0.34 | 0.31 | 0.43 | 0.46 | 0.63 | 0.55 | 0.47 | 0.50 | 0.67 | 0.60 |
| s0607_1_3 | 14.6 | 0.63 | 0.45 | 0.35 | 0.60 | 0.54 | 0.56 | 0.49 | 0.43 | 0.49 | **0.70** | **0.64** |
| s6687_1_4 | 15.4 | 0.49 | 0.42 | 0.19 | **0.63** | **0.53** | **0.57** | **0.58** | 0.40 | **0.55** | **0.51** | **0.53** |
| s0607_1_4 | 16.2 | 0.47 | 0.40 | 0.32 | **0.48** | 0.44 | 0.44 | **0.55** | 0.33 | 0.40 | **0.49** | **0.52** |
| s2687_1_4 | 16.3 | 0.61 | 0.28 | 0.22 | 0.51 | 0.43 | 0.53 | 0.56 | 0.35 | 0.45 | **0.65** | **0.67** |
| s0607_1_5 | 17.7 | 0.62 | 0.33 | 0.31 | 0.34 | 0.36 | 0.56 | 0.51 | 0.39 | 0.45 | **0.70** | **0.63** |
| s2307_1_1 | 19.9 | 0.64 | 0.27 | 0.30 | 0.34 | 0.36 | 0.50 | 0.50 | 0.49 | 0.50 | 0.62 | 0.55 |
| s1187_1_1 | 20.7 | 0.58 | 0.37 | 0.28 | **0.64** | 0.53 | 0.53 | 0.57 | 0.36 | 0.44 | nan | nan |
| s2307_1_2 | 21.7 | 0.60 | 0.27 | 0.29 | 0.31 | 0.29 | 0.45 | 0.50 | 0.37 | 0.41 | 0.58 | 0.50 |
| s8187_1_8 | 23.1 | 0.62 | 0.41 | 0.38 | 0.48 | 0.48 | 0.60 | 0.55 | 0.46 | 0.55 | **0.70** | 0.61 |
| Av. | 10.61 | 0.57 | 0.34 | 0.29 | 0.45 | 0.42 | 0.55 | 0.51 | 0.42 | 0.47 | 0.55 | 0.54 |

**Table S3. Average template and FreeSurfer data.** Hippocampal Dice metrics for each participant from: average linear/nonlinear infant template predicting CE and JF in linear/nonlinear space; average adult template predicting CE and JF in linear/nonlinear space; and FreeSurfer predicting CE and JF in native anatomical space. Bold indicates Dice metrics greater than IRR for that scan. "nan" denotes the two scans FreeSurfer failed to segment. Averages are presented in the final row.

| ID | Age | IRR | Adult-Pretrained-ASHS | | CE-ASHS | | JF-ASHS | | Infant-Trained-ASHS | |
|---|---|---|---|---|---|---|---|---|---|---|
| | | | CE | JF | CE | JF | CE | JF | CE | JF |
| s6057_1_1 | 3.6 | 0.50 | **0.60** | **0.56** | **0.66** | 0.47 | **0.53** | **0.64** | **0.62** | **0.59** |
| s6607_1_1 | 4.0 | 0.59 | **0.66** | **0.63** | **0.70** | **0.63** | 0.58 | **0.69** | **0.68** | **0.68** |
| s7067_1_1 | 4.7 | 0.46 | **0.53** | **0.49** | **0.59** | **0.50** | **0.51** | **0.58** | **0.56** | **0.54** |
| s6687_1_1 | 5.0 | 0.49 | **0.54** | **0.61** | **0.62** | **0.59** | 0.43 | **0.71** | **0.54** | **0.63** |
| s7017_1_1 | 5.2 | 0.48 | **0.70** | **0.53** | **0.77** | **0.50** | **0.54** | **0.69** | **0.68** | **0.62** |
| s0687_1_1 | 5.3 | 0.49 | **0.64** | **0.54** | **0.74** | **0.49** | **0.57** | **0.62** | **0.66** | **0.60** |
| s8687_1_2 | 5.8 | 0.57 | 0.51 | 0.57 | **0.64** | **0.62** | 0.46 | **0.60** | **0.60** | **0.65** |
| s2687_1_2 | 6.0 | 0.59 | **0.64** | **0.68** | **0.73** | **0.65** | 0.49 | **0.73** | **0.59** | **0.72** |
| s2037_1_2 | 6.5 | 0.64 | **0.73** | **0.66** | **0.75** | 0.59 | **0.66** | **0.73** | **0.74** | **0.69** |
| s7017_1_2 | 6.6 | 0.51 | **0.72** | **0.57** | **0.79** | 0.46 | **0.59** | **0.71** | **0.73** | **0.59** |
| s4607_1_2 | 6.7 | 0.49 | **0.57** | **0.61** | **0.62** | **0.64** | 0.44 | **0.68** | **0.54** | **0.70** |
| s1057_1_1 | 6.8 | 0.58 | **0.70** | 0.58 | **0.71** | 0.54 | **0.59** | **0.69** | **0.69** | **0.62** |
| s6607_1_2 | 7.4 | 0.63 | **0.68** | **0.69** | **0.69** | **0.64** | **0.64** | **0.75** | **0.67** | **0.73** |
| s3607_1_1 | 7.5 | 0.59 | **0.66** | **0.64** | **0.71** | **0.59** | **0.60** | **0.74** | **0.69** | **0.68** |
| s0607_1_1 | 7.6 | 0.68 | **0.74** | **0.71** | **0.75** | 0.63 | 0.66 | **0.75** | **0.74** | **0.72** |
| s6057_1_2 | 7.6 | 0.51 | **0.57** | **0.56** | **0.69** | **0.51** | **0.52** | **0.71** | **0.60** | **0.67** |
| s7017_1_4 | 8.3 | 0.50 | **0.60** | **0.57** | **0.70** | **0.52** | **0.50** | **0.61** | **0.61** | **0.61** |
| s8607_1_1 | 8.5 | 0.65 | **0.69** | **0.70** | **0.70** | 0.62 | 0.62 | **0.77** | **0.67** | **0.73** |
| s4107_1_1 | 8.8 | 0.53 | **0.77** | **0.59** | **0.76** | 0.53 | **0.62** | **0.73** | **0.73** | **0.63** |
| s0687_1_2 | 9.0 | 0.63 | **0.71** | **0.67** | **0.73** | 0.60 | **0.65** | **0.78** | **0.71** | **0.75** |
| s0057_1_3 | 9.0 | 0.56 | **0.69** | **0.60** | **0.70** | 0.52 | **0.65** | **0.72** | **0.70** | **0.67** |
| s0307_1_2 | 9.1 | 0.59 | **0.72** | 0.56 | **0.76** | **0.60** | **0.60** | **0.67** | **0.69** | **0.66** |
| s2687_1_3 | 9.9 | 0.60 | **0.62** | 0.60 | **0.71** | **0.63** | 0.51 | **0.68** | **0.62** | **0.66** |
| s1017_1_1 | 10.0 | 0.43 | **0.56** | **0.48** | **0.63** | **0.49** | **0.52** | **0.60** | **0.63** | **0.52** |
| s3607_1_2 | 10.4 | 0.50 | **0.59** | **0.59** | **0.70** | **0.56** | **0.53** | **0.72** | **0.63** | **0.66** |
| s1607_1_2 | 10.7 | 0.53 | **0.72** | **0.57** | **0.73** | 0.48 | **0.64** | **0.67** | **0.70** | **0.64** |
| s6607_1_3 | 10.8 | 0.66 | **0.76** | 0.65 | **0.77** | 0.63 | **0.70** | **0.75** | **0.77** | **0.71** |
| s6687_1_3 | 11.3 | 0.60 | **0.69** | **0.65** | **0.75** | 0.58 | 0.58 | **0.74** | **0.64** | **0.71** |
| s0607_1_2 | 12.7 | 0.58 | 0.56 | **0.62** | **0.67** | **0.59** | 0.56 | **0.72** | **0.64** | **0.69** |
| s4607_1_4 | 13.0 | 0.67 | **0.75** | **0.69** | **0.81** | 0.65 | 0.65 | **0.78** | **0.75** | **0.75** |
| s8187_1_4 | 13.8 | 0.65 | **0.77** | 0.64 | **0.81** | 0.59 | **0.66** | **0.78** | **0.73** | **0.73** |
| s4607_1_5 | 14.1 | 0.62 | **0.76** | 0.61 | **0.79** | 0.61 | **0.64** | **0.75** | **0.69** | **0.72** |
| s8687_1_4 | 14.5 | 0.70 | **0.75** | 0.65 | **0.78** | 0.63 | 0.67 | **0.76** | **0.78** | **0.71** |
| s0607_1_3 | 14.6 | 0.63 | **0.77** | **0.65** | **0.79** | 0.61 | **0.66** | **0.78** | **0.75** | **0.73** |
| s6687_1_4 | 15.4 | 0.49 | **0.69** | **0.61** | **0.77** | **0.57** | **0.56** | **0.76** | **0.71** | **0.67** |
| s0607_1_4 | 16.2 | 0.47 | **0.54** | **0.55** | **0.62** | **0.53** | **0.54** | **0.66** | **0.60** | **0.61** |
| s2687_1_4 | 16.3 | 0.61 | **0.70** | **0.63** | **0.77** | 0.59 | 0.60 | **0.77** | **0.70** | **0.72** |
| s0607_1_5 | 17.7 | 0.62 | **0.78** | **0.65** | **0.82** | 0.61 | **0.67** | **0.80** | **0.76** | **0.72** |
| s2307_1_1 | 19.9 | 0.64 | **0.77** | **0.66** | **0.77** | 0.62 | 0.63 | **0.79** | **0.70** | **0.76** |
| s1187_1_1 | 20.7 | 0.58 | **0.77** | **0.63** | **0.78** | 0.58 | **0.62** | **0.76** | **0.68** | **0.72** |
| s2307_1_2 | 21.7 | 0.60 | **0.73** | **0.61** | **0.71** | 0.59 | **0.62** | **0.70** | **0.69** | **0.66** |
| s8187_1_8 | 23.1 | 0.62 | **0.74** | **0.67** | **0.77** | **0.62** | 0.61 | **0.75** | **0.71** | **0.70** |
| Av. | 10.61 | 0.57 | **0.68** | **0.61** | **0.73** | **0.58** | **0.59** | **0.72** | **0.67** | **0.67** |

**Table S4. Pre-trained and trained ASHS data.** Hippocampal Dice metrics for each participant from: Adult-Pretrained-ASHS predicting CE and JF; CE-ASHS predicting CE and JF; JF-ASHS predicting CE and JF; and Infant-Trained-ASHS predicting CE and JF. Bold indicates Dice metrics greater than IRR for that scan. Averages are presented in the final row.
